# Mechanistic insights into the enhancement or inhibition of phase separation by polyubiquitin chains of different lengths or linkages

**DOI:** 10.1101/2021.11.12.467822

**Authors:** Thuy P. Dao, Yiran Yang, Maria F. Presti, Michael S. Cosgrove, Jesse B. Hopkins, Weikang Ma, Stewart N. Loh, Carlos A. Castañeda

## Abstract

Ubiquitin-binding shuttle UBQLN2 mediates crosstalk between proteasomal degradation and autophagy, likely via interactions with K48- and K63-linked polyubiquitin chains, respectively. UBQLN2 is recruited to stress granules in cells and undergoes liquid-liquid phase separation (LLPS) *in vitro*. However, interactions with ubiquitin or multivalent K48-linked chains eliminate LLPS. Here, we found that, although some polyubiquitin chain types (K11-Ub4 and K48-Ub4) did generally inhibit UBQLN2 LLPS, others (K63-Ub4, M1-Ub4 and a designed tetrameric ubiquitin construct) significantly enhanced LLPS. Using nuclear magnetic resonance (NMR) spectroscopy and complementary biophysical techniques, we demonstrated that these opposing effects stem from differences in chain conformations, but not in affinities between chains and UBQLN2. Chains with extended conformations and increased accessibility to the ubiquitin binding surface significantly promoted UBQLN2 LLPS by enabling a switch between homotypically to partially heterotypically-driven phase separation. Our study provides mechanistic insights into how the structural and conformational properties of polyubiquitin chains contribute to heterotypic phase separation with ubiquitin-binding shuttles and adaptors.

**Highlights:**

- Ubiquitin or short polyubiquitin chains bind to phase separation-driving stickers on UBQLN2 and inhibit its phase separation whereas longer chains provide the multivalency needed to enhance UBQLN2 phase separation.
- Phase separation of UBQLN2 is promoted over a wide range of Ub:UBQLN2 ratios in the presence of extended M1- and K63-linked Ub4 chains, but not compact K11- and K48-linked Ub4 chains.
- Chain conformation and accessibility of the Ub interacting surface is a driving factor of UBQLN2/polyUb co-phase separation.
- UBQLN2 condensates assemble during *in vitro* enzymatic assembly of K63-linked polyUb chains as free ubiquitin is reduced.

## Introduction

Protein quality control (PQC) mechanisms, which enable cells to combat aberrant proteins, is essential for cellular functions and viability (Dikic, 2017; Vendruscolo, 2012). Several distinct but connected pathways, such as proteasomal degradation, endoplasmic reticulum-associated protein degradation and autophagy exist to ensure proper PQC. In most cases, ubiquitination of substrate proteins is the common signal for these different pathways, as well as numerous other processes, such as DNA damage response, cargo transport and cell cycle control (Komander and Rape, 2012). This wide range of cellular responses to ubiquitination stems from the ability of ubiquitin (Ub) to form polyubiquitin (polyUb) chains with different lengths and linkages. The two most abundant and well-studied polyUb chain types are K48- and K63-linked chains that generally signal for proteasomal degradation and autophagy, respectively. Other chains, such as K11- and M1-linked chains, are involved in degradative and non-proteolytic pathways, respectively (Akutsu et al., 2016). It is hypothesized that polyUb chains elicit different biological signaling outcomes based on differences in their conformational properties and how each chain type interacts with Ub-binding receptors, such as shuttle proteins (Komander and Rape, 2012). Different shuttle proteins can recognize specific chains and transport modified substrates to distinct pathways (Komander and Rape, 2012). Some shuttle proteins, such as autophagy receptor NBR1 and proteasomal shuttle protein hHR23B, are mostly pathway specific. Others, such as UBQLN2 and p62, are involved in multiple pathways (Zientara-Rytter and Subramani, 2019). How shuttle proteins choose a specific pathway upon binding to ubiquitinated substrates, possibly through changes in conformations or oligomeric states, is of great interest (Lu et al., 2017).

PQC impairment due to aging, prolonged stress or mutations can lead to protein-containing inclusions characteristic of neurodegenerative diseases, such as amyotrophic lateral sclerosis (ALS), Parkinson’s and Alzheimer’s diseases (Hipp et al., 2014; Labbadia and Morimoto, 2015). These inclusions often comprise Ub, polyUb chains, Ub-binding shuttle proteins and other PQC proteins (Lowe et al., 1988; Manetto et al., 1988; Morimoto et al., 2015; Riley et al., 2010). Many of these proteins are also part of membraneless compartments, such as stress granules and PQC compartments that sequester aberrant proteins in preparation for clearance in healthy eukaryotic cells (Sontag et al., 2017). Dysregulation of membraneless compartments can lead to disease-linked inclusions (Molliex et al., 2015; Nedelsky and Taylor, 2019; Patel et al., 2015; Ryan and Fawzi, 2019). Liquid-liquid phase separation (LLPS) of a few key scaffolding proteins drive the formation of membraneless compartments whereas interacting partners of scaffolding proteins can modulate the properties of these compartments (Guillén-Boixet et al., 2020; Sanders et al., 2020; Yang et al., 2020; Yasuda et al., 2020). Therefore, knowledge of how binding partners affect phase-separating scaffolds is essential for understanding the regulation of PQC mechanisms.

We recently showed that Ub-binding shuttle protein UBQLN2 phase separates under physiological conditions but specific interactions with Ub, K48-linked Ub2 and Ub4 disrupt UBQLN2 LLPS (Dao et al., 2018). Following the polyphasic linkage formalism put forth by Wyman and Gill, UBQLN2 is considered a scaffold that drives condensate formation and Ub/K48 polyUb chains are ligands that can modulate LLPS (Wyman and Gill, 1980). Ligands can either enhance or inhibit LLPS, depending on their multivalency (in this case, the number of Ub units in a chain) and binding preferences for the scaffold in the dense and dilute phases (Ruff et al., 2021a). Unlike UBQLN2, proteasomal shuttle protein hHR23B and autophagy receptor p62 require longer K48- and K63-linked polyUb chains, respectively, to phase separate and carry out their functions (Sun et al., 2018; Yasuda et al., 2020). Interestingly, UBQLN1, which is a close homolog of UBQLN2 and is involved in both autophagy and proteasomal degradation, might bind preferentially to K63-over K48-linked chains (Harman and Monteiro, 2019). Therefore, we hypothesized, and showed in this work, that different polyUb chain types elicit distinct effects on UBQLN2 LLPS. By studying the interactions between UBQLN2 and various naturally occurring and designed polyUb chains, we determined the molecular mechanisms underlying these drastically different effects on UBQLN2 LLPS. Our results offer insights into how polyUb chains of distinct linkages drive ligand-induced phase transitions, and how modulation of UBQLN2 LLPS may steer UBQLN2 involvement into various PQC pathways.

## Results

### PolyUb chains with distinct linkage types and lengths differentially affect UBQLN2 LLPS

To determine if various polyUb chains impact UBQLN2 LLPS behavior differently, we initially focused on enzymatically synthesized K48 and K63 chains of two, three and four Ub units (Fig. S1A). These two chains are the most abundant in cells, signal for different cellular events, exhibit distinct solution conformations, and selectively drive LLPS of other Ub-binding shuttle proteins (Castañeda et al., 2016a; Dao and Castañeda, 2020; Sun et al., 2018; Yasuda et al., 2020; Ye et al., 2012).

We first screened for the effects of polyUb chains on full-length (FL) UBQLN2 LLPS by imaging UBQLN2 droplets that settle onto the coverslip at different Ub:UBQLN2 ratios (Fig. 1A). When LLPS was first induced, droplets from different solutions appeared to be similar in size. However, samples with increased LLPS contained more droplets that subsequently fuse and settle as large droplets on the coverslip. Consistent with our previous data, increasing amounts of Ub disassembled UBQLN2 droplets (Dao et al., 2018, 2019). Using nuclear magnetic resonance (NMR) spectroscopy, we confirmed that Ub binds only to the Ub-associated (UBA) domain in UBQLN2 with a K_d_ of 3 μM (similar to K_d_ for Ub binding to UBQLN2 450-624 (Dao et al., 2018)); only amide peaks of UBA residues shifted in the presence of Ub (Fig. S2A). As Ub binds to the same UBA sites that are important for UBQLN2 LLPS (Fig. S2B, (Dao et al., 2018)), we classified Ub as a monovalent ligand that interacts with UBA stickers and destabilizes LLPS (Ruff et al., 2021a).

**Figure 1.**
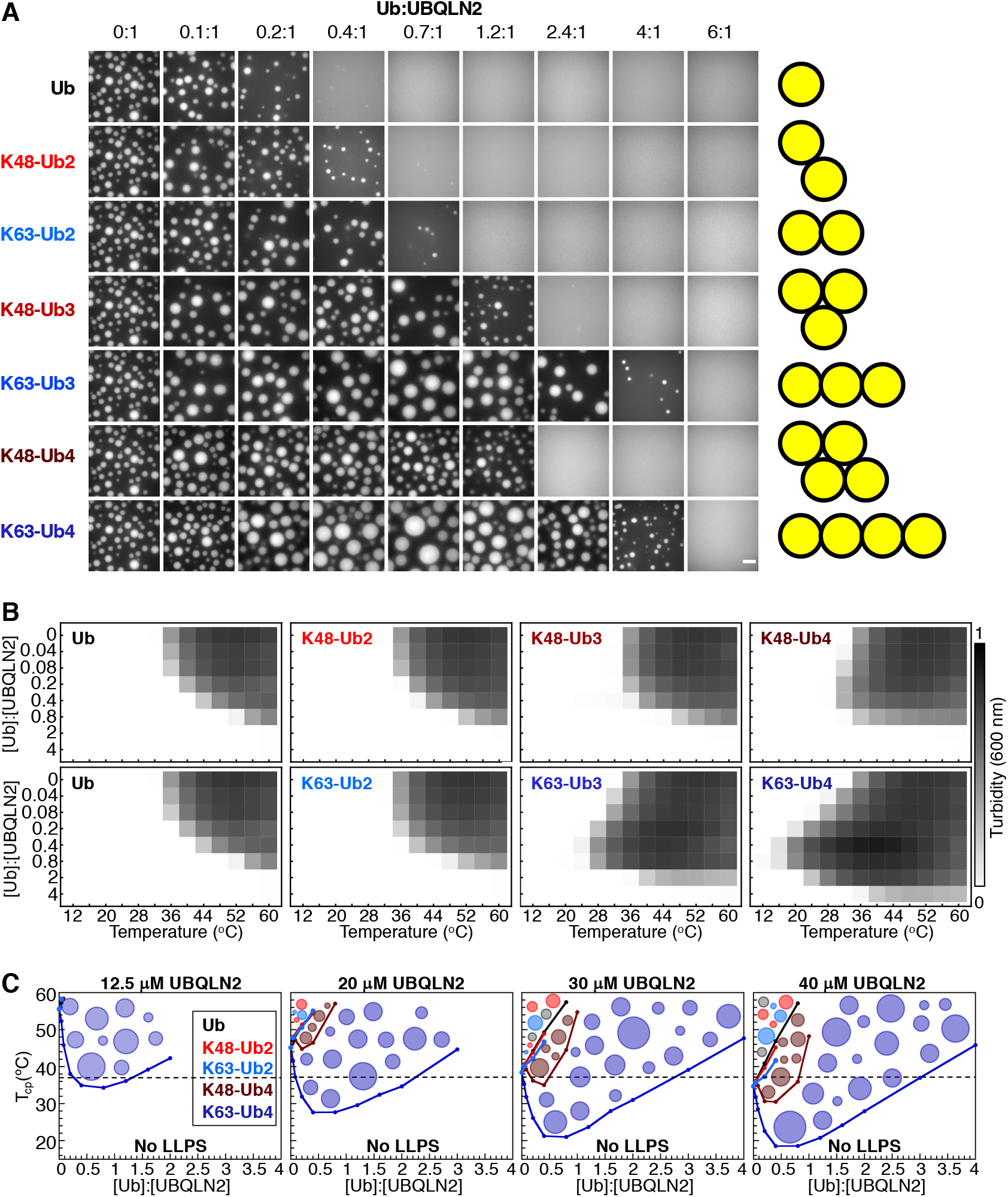
PolyUb chains with varied lengths and linkages differentially affect UBQLN2 LLPS. (A) Fluorescence microscopy showing UBQLN2 (Alexa Fluor 647) in solutions contained 50 μM FL UBQLN2 with increasing amounts of K48- and K63-linked chains of different lengths in 20 mM NaPhosphate and 200 mM NaCl (pH 6.8) at 30 °C. [Ub]:[UBQLN2] reflects the ratio between Ub monomers and UBQLN2 molecules. The same image is used for the 0:1 condition. Scale bar, 5 μm. Yellow cartoons illustrate one major solution conformation of each chain. (B) Representative results from spectrophotometric turbidity assay comparing the effects of varying amounts of K48- and K63-linked chains of different lengths on UBQLN2 LLPS at 30 μM UBQLN2. (C) Temperature–composition phase diagrams showing changes in cloud point temperatures as a function of Ub:UBQLN2. The black dashed lines denote biologically relevant temperature 37 °C, at which 12.5 μM UBQLN2 only undergoes LLPS with K63-Ub4.

In contrast to Ub, K48- and K63-Ub2, Ub3, and Ub4 chains can be considered di-, tri- and tetra-valent ligands that bind to UBQLN2. We observed several trends when mixing these multivalent ligands with UBQLN2: 1) Droplets were observed at increasingly higher Ub:UBQLN2 ratios as chain length/multivalency increased, regardless of linkage type. 2) Longer K63 chains significantly promoted UBQLN2 LLPS at lower Ub:UBQLN2 ratios. 3) All chains inhibited LLPS at higher ratios (Fig. 1A).

### K63-Ub4 promotes reentrant phase behavior of UBQLN2

To quantitatively describe the effects of polyUb on UBQLN2 LLPS, we ran turbidity experiments that monitored the change in A_600_ values as a function of temperature at different Ub:UBQLN2 ratios. The UBA domain of UBQLN2 was folded and able to interact with Ub at the temperature range used in our assays (Fig. S1C). We previously showed that high and low A_600_ values correlate with droplet formation and clearance, respectively, and that UBQLN2 phase separates with increasing temperature (Dao et al., 2018). Increasing amounts of Ub and K48-Ub2 reduced and eventually eliminated UBQLN2 LLPS (Fig. 1B and (Dao et al., 2018)). The turbidity plots for K63-Ub2 addition to UBQLN2 were nearly identical to those of K48-Ub2 and Ub, suggesting that lysine linkage does not matter at the Ub2 level.

Addition of Ub3 and Ub4 chains of different linkages led to distinct turbidity plots (Fig. 1B, Fig. S1B), consistent with microscopy data (Fig. 1A). Strikingly, K63-Ub4 visibly promoted and inhibited LLPS, depending on Ub:UBQLN2 ratio, reminiscent of reentrant phase behavior seen for protein-nucleic acid and protein-protein coacervates that form via heterotypic interactions (Alshareedah et al., 2019; Banerjee et al., 2017; Choi et al., 2019; Dignon et al., 2020; Feric et al., 2021; Xu et al., 2020). Low K63-Ub3/Ub4:UBQLN2 ratios significantly promoted LLPS, driving UBQLN2 droplet assembly at lower temperatures than in the absence of K63-Ub3/Ub4. As these ratios were further increased, LLPS was inhibited. We hypothesized that K63-Ub4 promotes LLPS by acting as an emergent scaffold onto which multiple UBQLN2 molecules bind. These heterotypic interactions provide additional multivalency to further promote LLPS. However, as Ub4 concentrations increase, not enough UBQLN2 molecules exist to bind Ub4 simultaneously, hence diluting out the LLPS-driving UBQLN2-UBQLN2 interactions and leading to inhibition of LLPS (Banerjee et al., 2017; Choi et al., 2019; Dao and Castañeda, 2020; Ruff et al., 2021b). Conversely, low K48-Ub3/Ub4:UBQLN2 ratios only slightly enhanced LLPS (Fig. S1B). Increasing addition of K48-Ub3/Ub4 inhibited LLPS at much lower Ub:UBQLN2 ratios than for K63-Ub3/Ub4.

To quantitatively compare the effects of polyUb chains on UBQLN2 LLPS, we obtained temperature-composition phase diagrams for different Ub:UBQLN2 ratios at multiple UBQLN2 concentrations (Fig. 1C). We determined T_cp_(infl), the cloud point temperature at the inflection point of the transition (Yang et al., 2019). We used T_cp_ at different Ub:UBQLN2 ratios to map out the coexistence curve, above which the protein solution is phase separated. For Ub and Ub2, we observed linear increase in T_cp_ as Ub:UBQLN2 ratio increased, consistent with Ub and Ub2 being low-valency ligands that interact with UBQLN2 UBA “stickers” (UBQLN2 residues that are important for self-association and LLPS) to drive droplet disassembly (Dao et al., 2018). In contrast, we observed a substantial decrease in T_cp_ at low K63-Ub4:UBQLN2 ratios, indicating a greater regime of the phase diagram over which UBQLN2 is phase separated. This effect was clearest at the lowest concentration tested (12.5 μM) that approaches the physiological UBQLN2 concentration (1-2 μM) (Dao et al., 2018). At 37 °C, 12.5 μM UBQLN2 did not phase separate by itself or with Ub, K48-Ub2, K63-Ub2, and K48-Ub4, but readily phase separated with K63-Ub4 between Ub:UBQLN2 ratios of 0.3:1 (~ 1 μM K63-Ub4) and 1.3:1 (~ 4 μM K63-Ub4). Our data illustrate a transition from UBQLN2 LLPS driven by homotypic interactions to a system where LLPS is driven, at least partially, by heterotypic interactions between UBQLN2 and K63-Ub4 (Dao and Castañeda, 2020). Therefore, polyUb chains of different linkages can substantially alter the driving forces of phase-separating systems.

### NMR and fluorescence anisotropy titrations reveal differences in how K48- and K63-Ub4 interact with UBQLN2

To examine the molecular origins of the differential effects of K48- or K63-Ub4 on UBQLN2 LLPS, we used NMR to map the interactions between Ub4 and UBQLN2. First, we looked at FL UBQLN2 in the presence of Ub4, but the UBA resonances were severely attenuated, precluding quantitative NMR studies (Fig. S3A). To circumvent this issue, we used UBQLN2 450-624, a C-terminal construct that exhibits LLPS behavior similar to FL UBQLN2 and is amenable to NMR studies (Dao et al., 2018, 2019). We obtained temperature-composition phase diagrams using UBQLN2 450-624 and different amounts of K48- or K63-Ub4. Our phase diagrams showed that the two Ub4s alter LLPS of 450-624 in a similar manner as for FL UBQLN2 (Fig. S3B). Therefore, for subsequent NMR experiments, we collected data using UBQLN2 450-624 under non-phase separating conditions.

Upon titration of K48- or K63-Ub4 into ^15^N-labeled UBQLN2 450-624, we observed broadening of UBA peaks, likely due to the large size of the Ub4:UBQLN2 450-624 complex (> 35 kDa Ub4 + 1-4X 17 kDa UBQLN2 450-624). Despite this, we could follow the resonances and observe chemical shift perturbations (CSPs) mainly for UBA residues (Fig. 2A), indicating that Ub units in the chains bind primarily to UBA and in a manner like monoUb. However, we also observed CSPs (Fig. 2A) and reduced peak intensity (Fig. 2B) for residues 555-570, more so with K48-Ub4 than K63-Ub4, which adopt compact and extended solution conformations in the unbound state, respectively (Tenno et al., 2004). We recapitulated the same trends with UBQLN2 487-624 (Fig. 2C). Resonances in UBQLN2 487-624 exhibit better NMR signal to noise quality since this construct does not oligomerize, undergo LLPS, or exhibit backbone dynamics on a slow millisecond timescale (Dao et al., 2018). Residues 555-570 were not perturbed in the presence of monoUb (Dao et al., 2018). Furthermore, we did not observe CSPs outside of the UBA domain in a UBQLN2 450-624 construct lacking residues 551-569 (Fig. 2D). Residues 555-570, but not the surrounding regions, exhibit slight helical propensity (Dao et al., 2018). Together, these data suggest that the transiently helical 555-570 region either specifically interacts with K48-Ub4 or is sterically affected when the compact K48-Ub4 binds to UBA. Since the 555-570 region is a “sticker” that mediates UBQLN2 LLPS (Dao et al., 2018), we speculated that K48-Ub4 may inhibit UBQLN2 LLPS by preventing both UBA and 555-570 stickers from contributing to LLPS.

**Figure 2.**
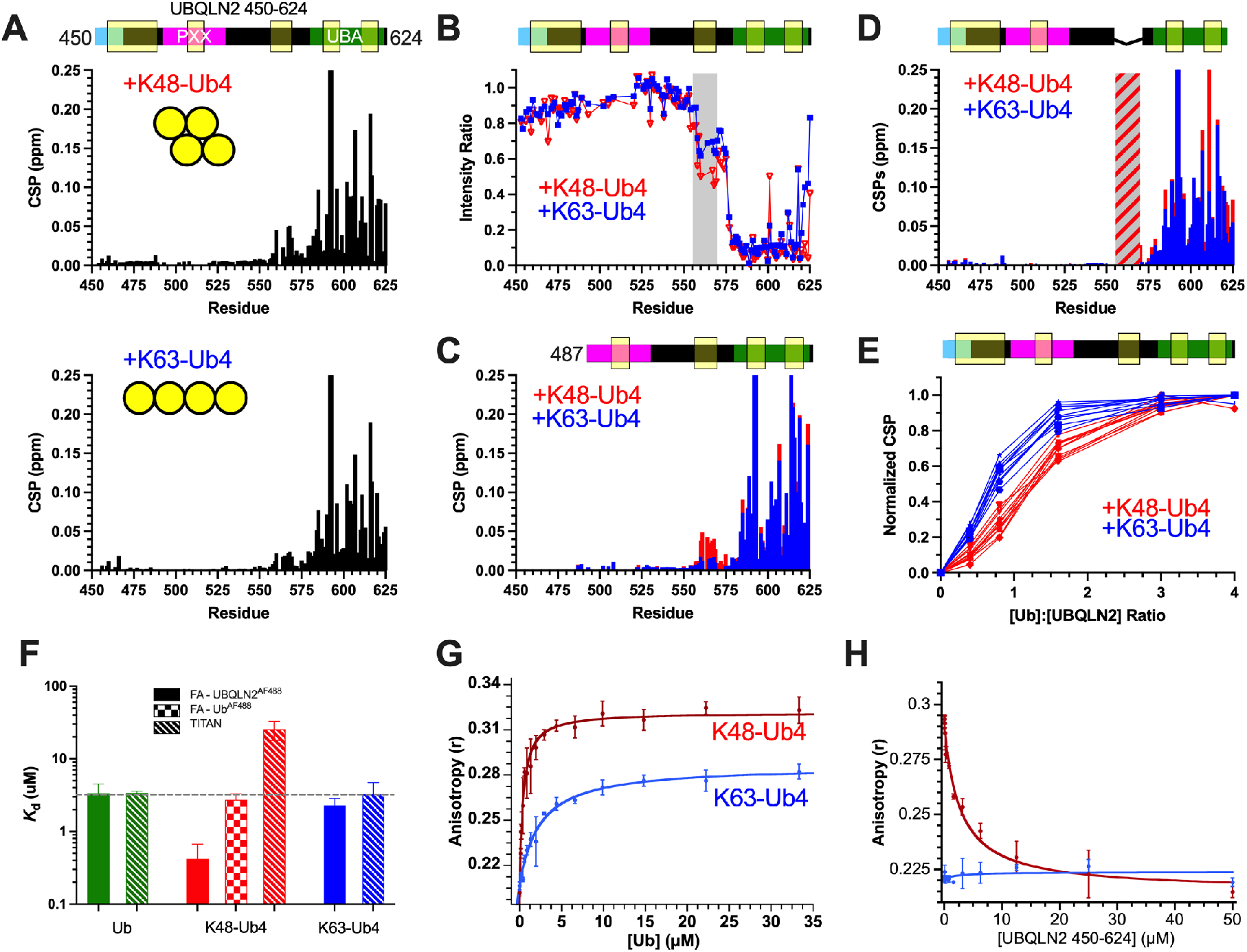
K48- and K63-Ub4 bind differently to UBQLN2 450-624. (A) CSPs for UBQLN2 450-624 residues at 1:1 ratio with Ub4. The highlighted regions in the domain map are important for the homotypic interactions that drive UBQLN2 LLPS (Dao et al., 2018). (B) Amide peak intensities between the unbound and Ub4-bound forms of UBQLN2 450-624 with the 555-570 region highlighted in gray. (C) CSPs for residues in UBQLN2 487-624 at 1:1 ratio with Ub4. (D) CSPs for residues in UBQLN2 450-624 lacking residues 551-569 at 1:1 ratio with Ub4. (E) Normalized titration curves of 11 amide resonances for UBQLN2 450-624. (F) Comparison of binding affinity (K_d_) measurements obtained from NMR and fluorescence anisotropy. (G) Fluorescence anisotropy values for fluorescently-labeled UBQLN2 450-624 in the presence of increasing K48- or K63-Ub4. (H) Fluorescence anisotropy values for fluorescently-labeled K48- or K63-Ub4 in the presence of increasing UBQLN2 450-624.

To determine the binding affinities between K48- or K63-Ub4 and UBQLN2, we collected residue-specific titration curves for resonances in UBQLN2 450-624 with the two Ub4s (Fig. 2E). The curves for UBQLN2 amide resonances were sigmoidal, especially for the titration with K48-Ub4, and resonances exhibited extensive line broadening. These conditions preclude us from considering CSPs as simple weighted averages of free and bound species, as we are no longer under fast exchange conditions (Williamson, 2013). Instead, we used TITAN 2D lineshape analysis (Waudby et al., 2016), which accounts for both CSP and peak broadening during the titration, to obtain K_d_ values. Assuming single-site binding between UBQLN2 and each Ub unit in Ub4, we obtained binding affinities of 3 μM for both Ub and K63-Ub4, and a lower affinity of 25 μM for K48-Ub4 (Fig. 2F, Fig. S6, Table S1). We corroborated our NMR measurements of K_d_ values using fluorescence anisotropy experiments. Anisotropy monitors how quickly the fluorophore tumbles and reorients in solution, which are influenced by the size and flexibility of the protein to which the fluorophore is attached (higher anisotropy indicates larger size and/or decreased flexibility). We observed increases in anisotropy on titrating K48-Ub4, K63-Ub4, and Ub into fluorescently-labeled UBQLN2 450-624 (Fig. 2F, 2G). The fitted K_d_ values for UBQLN2 with Ub, K48-Ub4 and K63-Ub4 were 3.4, 0.4 and 2.3 μM, respectively (Fig. 2F, Table S1). Interestingly, like NMR, anisotropy reported very similar K_d_ values for binding between UBQLN2 and Ub or K63-Ub4, but not K48-Ub4.

To further investigate the differences in K_d_ values for K48-Ub4 between NMR and anisotropy, we carried out the reciprocal anisotropy experiments in which we titrated UBQLN2 450-624 into K48-Ub4 or K63-Ub4 containing a fluorescently labeled Ub unit (see Methods). There was minimal change in anisotropy for K63-Ub4 with increasing amounts of UBQLN2, indicating minimal differences in how the complex tumbled compared to how the fluorescently labeled Ub unit in K63-Ub4 tumbled (Fig. 2H). In contrast, anisotropy significantly decreased upon K48-Ub4 binding to UBQLN2. This decrease in anisotropy is atypical and indicates that binding of UBQLN2 to K48-Ub4 caused the latter to become more flexible despite the increased mass of the complex (Fig. 2H). Titrating UBQLN2 into K48-Ub4 with a labeled Ub unit yielded a K_d_ of 2.7 μM, similar to the K_d_ values for UBQLN2 and K63-Ub4 by NMR and anisotropy. We hypothesized that the conflicting binding data from NMR and anisotropy stemmed from differences in conformations of K48-Ub4 and K63-Ub4.

### The Ub-Ub interface in K48-Ub4 “opens” to accommodate UBQLN2 binding

K48- and K63-Ub4 are known to adopt compact and extended conformations, respectively, with the latter containing Ub units in a beads-on-a-string arrangement (Fig. 3A). Indeed, small angle x-ray scattering (SAXS) (Fig. 4B, Table S2) showed that unbound K48-Ub4 is compact (R_g_ = 25.67 ± 0.05 Å), whereas K63-Ub4 is extended (R_g_ = 32.33 ± 0.09 Å), consistent with prior structural studies (Tenno et al., 2004; Varadan et al., 2002). Sedimentation velocity analytical ultracentrifugation (SV-AUC) data support these conclusions, as K48-Ub4 has a higher sedimentation coefficient and lower frictional ratio than K63-Ub4 (Fig. S4, Table S3). We also observed higher anisotropy for fluorescently-labeled UBQLN2 upon titration of K48-Ub4 compared to K63-Ub4 (Fig. 2G). These data suggest that, as the limiting reagent, only one UBQLN2 would bind to each K48-Ub4, which remains compact, and therefore, high anisotropy is observed for the large UBQLN2/K48-Ub4 complex. In contrast, when UBQLN2 is bound to an individual Ub bead on a K63-Ub4 string, the anisotropy for the UBQLN2/K63-Ub4 complex is not as high due to the smaller size of the tumbling unit. With the reciprocal titration where K48-Ub4 is the limiting reagent, anisotropy significantly decreased, suggesting that K48-Ub4 must open to accommodate simultaneous binding of multiple UBQLN2, leading to the fluorescently-labeled Ub tumbling more independently as a smaller unit from the rest of the Ub4 chain.

**Figure 3.**
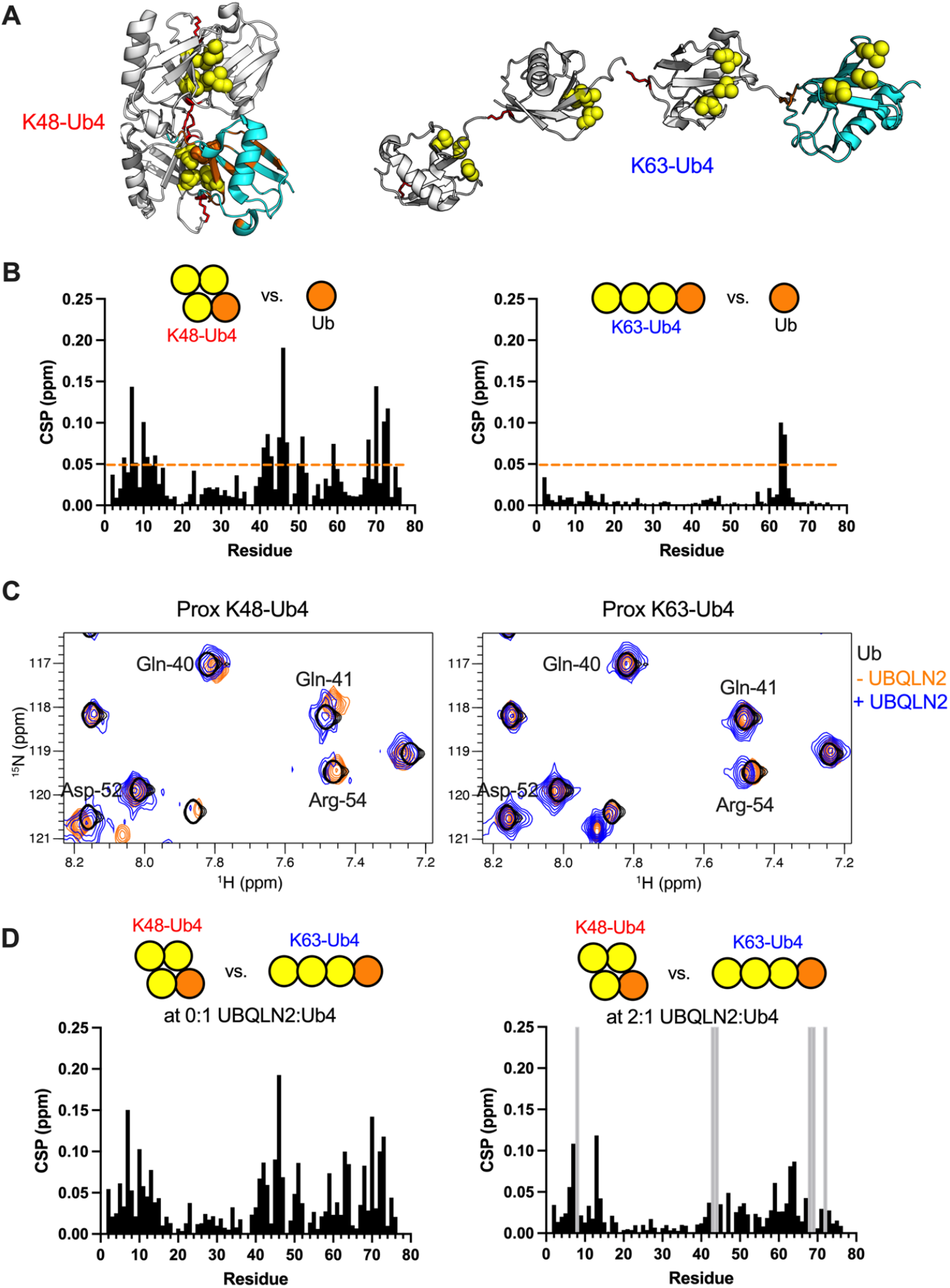
The Ub-Ub interface in K48-Ub4 opens to bind to UBQLN2. (A) Representative structures of K48-Ub4 (PDB ID 2O6V) and K63-Ub4 (PDB ID 3HM3). The cyan proximal Ub unit was selectively ^15^N-labeled for NMR binding experiments. Yellow spheres represent hydrophobic patch residues L8, I44, and V70. Red sticks indicate K48 or K63 linkage. CSPs > 0.05 ppm for proximal Ub resonances vs. monoUb positions (in the absence of UBQLN2) are color-coded orange. (B) Proximal Ub resonances in Ub4 were compared to monoUb, as noted in schematic. (C) ^15^N-^1^H SOFAST-HMQC NMR spectra of proximal Ub of K48-Ub4 or K63-Ub4 at 50 μM in the absence and presence of UBQLN2 450-624 at a 2:1 UBQLN2:Ub4 ratio. Unbound monoUb spectrum (black) is overlaid for comparison. NMR spectra collected and processed under identical conditions. (D) CSPs of the same proximal Ub resonances in K63-Ub4 vs K48-Ub4. Gray bars indicate K48-Ub4 resonances that were too weak to be detected.

**Figure 4.**
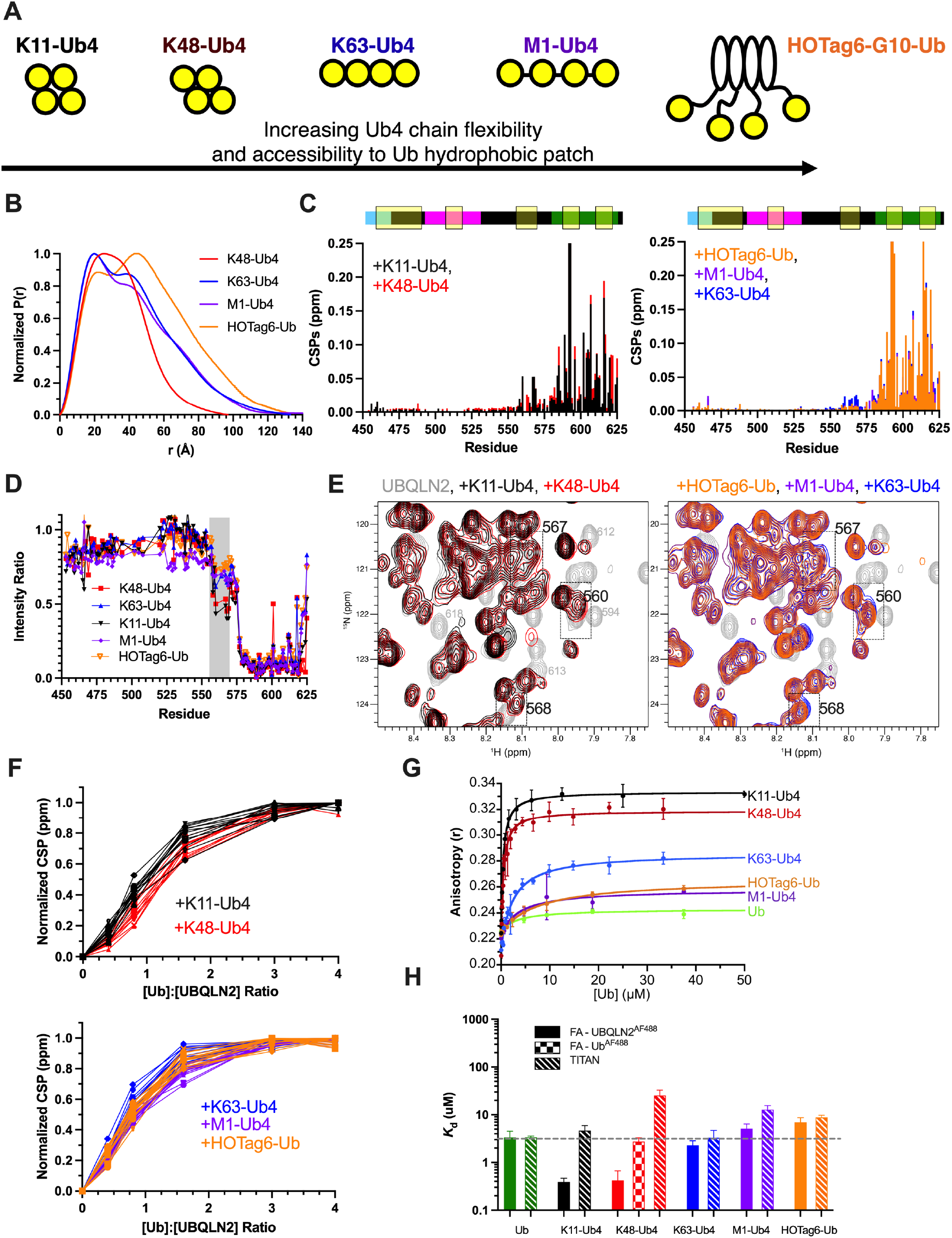
Compact K11 and K48-Ub4 bind UBQLN2 differently than extended K63-Ub4, M1-Ub4 and HOTag6-G10-Ub. (A) Cartoons depicting a possible solution conformation for each of the Ub4 chains studied here. (B) Normalized pair distance distribution function P(r) from SAXS data for four Ub4 chain types. (C) CSPs at 1:1 Ub4:UBQLN2 for all chain types, but clustered into two categories for clarity. (D) Peak intensity comparison in the absence and presence of Ub4 at 1:1 Ub4:UBQLN2. (E) TROSY-HSQC spectra of UBQLN2 450-624 in the absence and presence of different Ub4 chains. Some of the peaks in the 555-570 region are more perturbed in the presence of K11-Ub4 and K48-Ub4 but not the other chains, whereas the UBA peaks (580-624) are equally perturbed in the presence of any of the chains. (F) Normalized titration curves of UBA resonances showing more pronounced sigmoidal behaviors upon titration of K48-Ub4 into UBQLN2 450-624 than upon titrations of K11-Ub4, K63-Ub4, M1-Ub4 and HOTag6-Ub. (G) Fluorescence anisotropy values for fluorescently-labeled UBQLN2 450-624 in the presence of increasing amounts of different Ub4 chains. (H) Comparison of K_d_ values obtained from NMR and fluorescence anisotropy.

To test our hypothesis that K48-Ub4 changes from a compact to a more extended conformation upon interacting with UBQLN2 450-624, we performed NMR titration of UBQLN2 450-624 into either K48 or K63-Ub4 with ^15^N-labeled proximal Ub (Fig. 3A). First, we compared the HSQC spectrum of each proximal Ub unit to monoUb (in the absence of UBQLN2) and found significant CSPs for resonances in the proximal Ub of K48-Ub4, but not K63-Ub4, indicative of Ub-Ub interactions among K48-Ub4 subunits (Fig. 3B). The large CSPs map to the UBQLN2-binding hydrophobic patch of Ub (L8, I44, V70) and surrounding residues, consistent with known structures of K48-Ub4 (Fig. 3A, Eddins et al., 2007). Together with SAXS and AUC data (Figs. 4B and S4, Tables S2 and S3), these data suggested that the K48-Ub4 is compact in solution and that the Ub hydrophobic patches are not immediately available to bind ligands.

Upon titration of UBQLN2 450-624, we observed that several resonances near the Ub-Ub interface (residues 10, 39, 40, 41, 51, 52, 73, 75) in the ^15^N-labeled proximal Ub of K48-Ub4 converged to the locations of the same resonances in the unbound monoUb (Fig. 3C), indicating that these resonances in proximal Ub of K48-Ub4, as UBQLN2 was titrated, were in similar chemical environments as free monoUb. In contrast, the same resonances in proximal Ub of K63-Ub4, in which no Ub units are in close contact with another, did not move significantly with initial addition of UBQLN2 (Fig. 3C). In other words, the hydrophobic patch of the proximal Ub of K48-Ub4 exhibited different chemical environments than that of Ub and the proximal Ub of K63-Ub4 in the absence of UBQLN2 but was in similar local environments in the presence of UBQLN2 (Fig. 3D). These data are consistent with our hypothesis that K48-Ub4 needs to “open” to accommodate UBA binding, partially explaining the sigmoidal titration curves and the weaker NMR-derived apparent binding affinity for K48-Ub4 from the UBQLN2 450-624 side (see above and Table S1). Therefore, the Ub binding patch is readily accessible in K63-Ub4 but requires an additional conformational change in K48-Ub4 to interact with UBQLN2. These observations resulted from differences in chain conformations and not binding ability as SV-AUC experiments of K48- or K63-Ub4 in the presence of UBQLN2 487-624 at a 2:1 UBQLN2:Ub4 loading ratio showed complete binding between either chain with UBQLN2 (Fig. S4). The AUC data are consistent with a model where UBQLN2:K48-Ub4 complex is slightly more compact than the UBQLN2:K63-Ub4 complex (Table S3).

In summary, we suspect that K48-Ub4 does not readily promote UBQLN2 LLPS through heterotypic interactions because K48-Ub4 (1) interacts with the 555-570 “sticker” required for LLPS, (2) is compact in conformation, and (3) contains Ub hydrophobic patches that are not as readily accessible for UBQLN2 binding. In contrast, K63-Ub4 can act as a scaffold to promote LLPS since K63-Ub4 (1) minimally interacts with 555-570, (2) adopts an extended conformation, and (3) contains Ub hydrophobic patches that are readily accessible for UBQLN2 binding.

### Effects of K11, M1-Ub4 and tetrameric HOTag6-G10-Ub on UBQLN2 binding

From the K48 and K63-Ub4 data, we hypothesized that differences in Ub4 conformation and Ub “sticker” accessibility are important determinants that drive differential effects on LLPS of UBQLN2. To test this hypothesis, we extended our study to examine how UBQLN2 interacts with other naturally occurring Ub4 chains, specifically K11 and M1 linkages, and a designed tetrameric HOTag6-G10-Ub (Fig. 4A). K11-Ub4 exhibits a compact conformation similar to K48-Ub4, whereas M1-Ub4 is similar to K63-Ub4 in that it adopts an extended conformation and the Ub binding patches are readily accessible, as confirmed by SAXS (Fig. 4B, Fig. S5A). HOTag6-G10-Ub comprises HOTag6, which exists as a tetramer (Zhang et al., 2018), and tethered Ub, linked by 10 glycines, to form a tetrameric complex with high conformational flexibility between the Ub units (Fig. 4B, Fig. S5). Therefore, these three constructs, together with K48- and K63-Ub4, form a library of Ub tetramers with varying conformations, flexibility, and hydrophobic patch accessibility to UBQLN2 (Fig. 4A, 4B).

We used NMR spectroscopy to probe how these different polyUb chains interact with UBQLN2 450-624. As expected, we observed the largest CSPs for UBA resonances (Fig. 4C). CSPs for K11-Ub4 were very similar to those for K48-Ub4, whereas CSPs for M1-Ub4 and HOTag6-G10-Ub were very similar to those for K63-Ub4. Amide resonances for residues 555-570 were more perturbed and peak intensities for residues 555-570 were more reduced in the presence of either compact K11- or K48-Ub4 chains, but not with the extended or more conformationally flexible M1-Ub4, K63-Ub4 or HOTag6-G10-Ub (Fig. 4C–4E).

Using NMR titration experiments and TITAN 2D lineshape analysis, we measured binding affinities between labeled UBQLN2 450-624 and these chain types (Fig. 4F, Fig. S6). We obtained K_d_ values of 4.6, 12.8 and 8.8 μM for K11-Ub4, M1-Ub4, and HOTag6-G10-Ub, respectively (Fig. 4H, Table S1). Unlike K48-Ub4 which is compact with occluded Ub hydrophobic patches, K11-Ub4 is compact with more exposed Ub hydrophobic patches (Fig. S7). Correspondingly, we observed that UBQLN2 NMR titration curves with K11-Ub4 barely exhibit sigmoidal behavior as K11-Ub4 might not need to open to accommodate UBQLN2 binding (unlike that of K48-Ub4). Despite K11-Ub4, M1-Ub4, and HOTag6-G10-Ub exhibiting different conformations in solution, they exhibit similar K_d_ values with UBQLN2, indicative of minimal linkage preference between UBQLN2 and polyUb.

Fluorescence anisotropy experiments using fluorescently-labeled UBQLN2 450-624 revealed differences in the tumbling properties of the UBQLN2:Ub4 complexes (Fig. 4G). Upon addition of either K11- or K48-Ub4, the final anisotropy values for UBQLN2 were similarly high, suggesting that UBQLN2:K11-Ub4 and UBQLN2:K48-Ub4 complexes tumble slowly as large units. In contrast, the final anisotropy values for UBQLN2 were low with either HOTag6-G10-Ub or M1-Ub4, similar to the anisotropy value for UBQLN2 when saturated with Ub. Since both M1-Ub4 and HOTag6-G10-Ub comprise Ub units that are connected by highly flexible linkers (Fig. 4A and Fig. S7), these data suggest that the Ub units in these two proteins move relatively independently. The anisotropy-derived K_d_ values for M1-Ub4 and HOTag6-G10-Ub were 5.1 and 7.0 μM, respectively, similar to those determined by NMR and for K63-Ub4 and Ub.

We note a tighter apparent anisotropy-derived K_d_ for K11-Ub4 of 0.39 μM, similar to that of K48-Ub4 (Fig. 4H). Both K11- and K48-Ub4 adopt compact conformations but have differently accessible UBQLN2-binding sites, possibly leading to discrepancies between NMR- and anisotropy-derived K_d_ values. As UBQLN2 reagent is limiting (100 nM) under the anisotropy experimental conditions, the UBQLN2:Ub4 complexes likely only have a single UBQLN2 bound. Given the compact arrangements of Ub units in both K11- and K48-Ub4, we speculate that when the UBQLN2 UBA domain binds to a Ub unit, a secondary interaction occurs between UBQLN2 and the rest of K11- or K48-Ub4. Indeed, we observed increased UBQLN2 CSPs for residues 555-570 adjacent to the UBA domain for both K11- and K48-Ub4 (Fig. 4C), although these CSPs are significantly weaker than those observed for the UBA domain.

Together, our data suggest that all five chains have similar binding affinities to UBQLN2. However, because of their quaternary structures and solution conformations, we predicted that they may have different effects on UBQLN2 LLPS. Specifically, K11-Ub4 may affect UBQLN2 LLPS similarly to K48-Ub4 due to CSPs for residues 555-570 and K11-Ub4’s compact conformation, whereas M1-Ub4 and HOTag6-G10-Ub may affect UBQLN2 LLPS similarly to K63-Ub4 due to similarities in binding affinity, extended conformation, and readily accessible Ub binding patches.

### Extended and flexible, but not compact, polyUb chains promote UBQLN2 LLPS

Next, we performed microscopy and turbidity assay experiments to determine how K11-Ub4, M1-Ub4 and HOTag6-G10-Ub affect UBQLN2 LLPS. Microscopy showed that M1-Ub4 and HOTag6-G10-Ub stabilized UBQLN2 droplets over a wide range of Ub:UBQLN2 ratios (Fig. 5A), as confirmed by turbidity plots (Fig. 5B) and temperature-concentration phase diagrams (Fig. 5C). Like K48-Ub4, addition of K11-Ub4 largely disassembled UBQLN2 droplets. Strikingly, M1-Ub4 and HOTag6-G10-Ub stabilized UBQLN2 LLPS over a larger Ub:UBQLN2 range than K63-Ub4. Importantly, reentrant phase behavior, by which high concentrations of polyUb chains disassemble UBQLN2 droplets, still existed for all chains. As flexibility and propensity to adopt an extended conformation increased for polyUb (HOTag6-Ub > M1-Ub4 > K63-Ub4 > K48 or K11-Ub4), UBQLN2/polyUb LLPS was stabilized and promoted. These data bolster our hypothesis that more extended and flexible chains with more accessible Ub binding patches can promote UBQLN2 LLPS to a greater extent.

**Figure 5.**
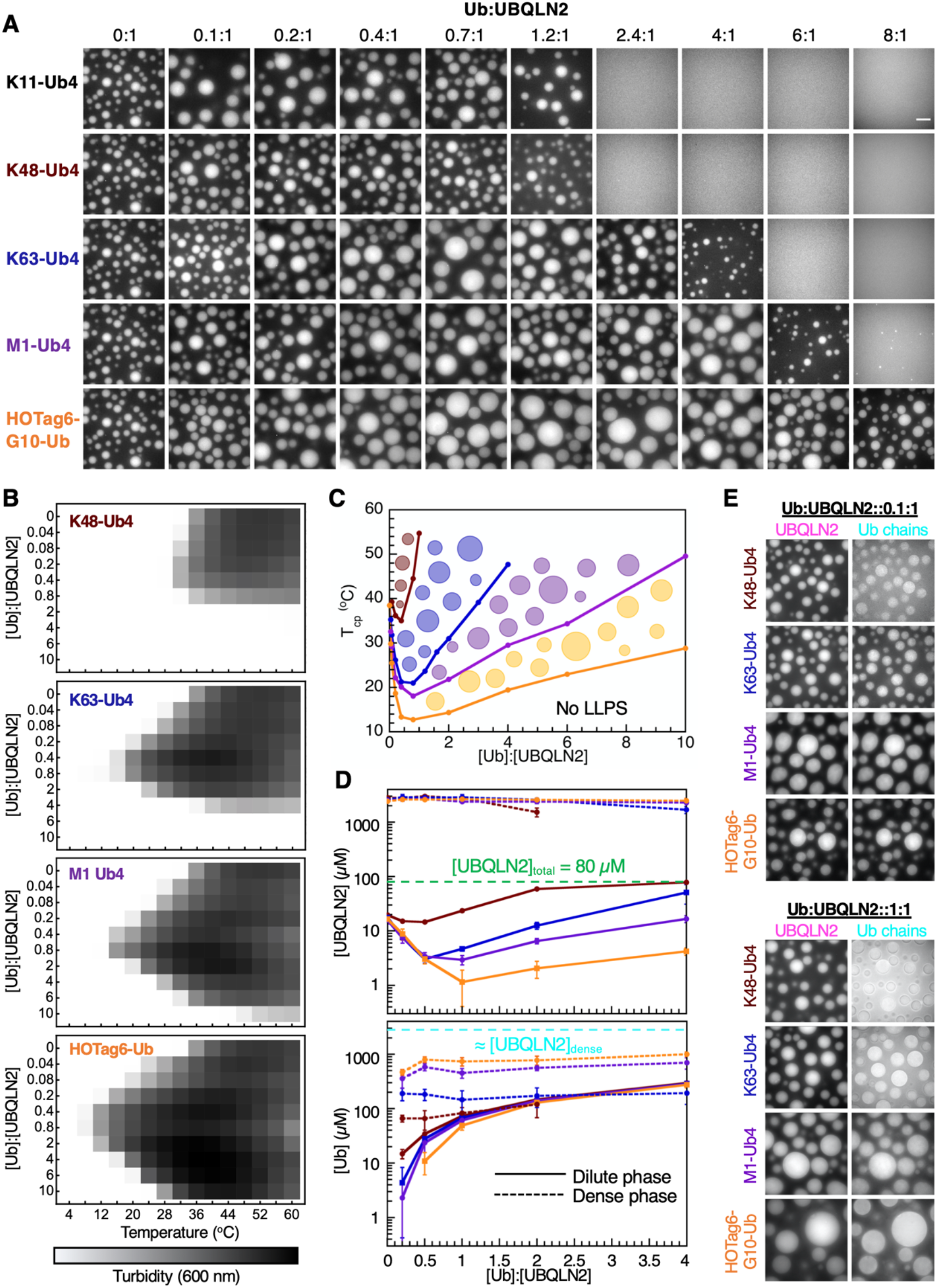
The effects of polyUb chains on UBQLN2 LLPS are dependent on chain flexibility / accessibility to Ub binding patch. (A) Fluorescence microscopy showing enhancement or inhibition of UBQLN2 droplet formation at increasing amounts of chains at 50 μM UBQLN2 in 20 mM NaPhosphate and 200 mM NaCl (pH 6.8) at 30 °C. [Ub]:[UBQLN2] reflects the ratio between Ub monomers and UBQLN2 molecules. Solution is spiked with Alexa Fluor 647-labeled UBQLN2. The same image is used for the 0:1 condition. Scale bar, 5 μm. (B) Representative results from spectrophotometric turbidity assay as a function of temperature at 30 μM UBQLN2. (C) Temperature–component phase diagrams showing the changes in LCST phase transition cloud-point temperatures as a function of Ub:UBQLN2 for different Ub4 at 30 μM UBQLN2. (D) Phase diagrams showing the dense and dilute phase concentrations of UBQLN2 (top) and Ub chains (bottom). Error bars represent the standard deviation over three trials. (E) Fluorescence microscopy showing the distribution of Ub4 (spiked with Dylight 488-labeled chains) inside and outside droplets containing 50 μM UBQLN2 (spiked with Alexa Fluor 647-labeled UBQLN2).

To determine how polyUb affect UBQLN2 concentration in the dilute and dense phases, we labeled UBQLN2 and Ub4 with organic dyes that absorb at different wavelengths. We induced UBQLN2 LLPS with Ub4 at different Ub:UBQLN2 ratios, centrifuged to separate the two phases, and measured the absorbance of the two dyes to extract the dilute and dense phase concentrations of both UBQLN2 and Ub4 (see Methods, Fig. 5D). The UBQLN2 dilute phase concentration, or c_sat_, was entirely consistent with our phase diagrams (Fig. 5C). In the presence of K48-Ub4, UBQLN2 c_sat_ barely changed at low Ub:UBQLN2 ratios, but increased to total UBQLN2 concentration (only dilute phase existed) at high ratios. Conversely, at low Ub:UBQLN2 ratios for K63, M1-Ub4 and HOTag6-G10-Ub, c_sat_ significantly decreased before increasing again at high ratios. Effectively, the change in c_sat_ mirrored the UBQLN2/polyUb phase diagrams, in the order of increasing chain flexibility (Fig 4A). Intriguingly, although the addition of Ub4 significantly changed the UBQLN2 dilute phase concentration, the UBQLN2 dense phase concentration remained the same in the presence of any Ub4, regardless of Ub:UBQLN2 ratios. These data suggest that the addition of K63-Ub4, M1-Ub4 and HOTag6-G10-Ub chains regulates UBQLN2 LLPS by changing the volume fraction, but not concentration, of the dense phase, consistent with a previous computational study for generic systems (Ruff et al., 2021a, 2021b). Therefore, HOTag6-G10-Ub, and to lesser extents K63 and M1-Ub4, can enhance UBQLN2 LLPS without diluting the UBQLN2 dense phase concentration over a wide range of polyUb ligand concentrations.

Unlike for UBQLN2, c_sat_ for Ub4 increased with increasing Ub:UBQLN2 ratios or total Ub concentrations (Fig. 5D). At low ratios, c_sat_ for Ub4 were in the following order: K48-Ub4 > K63-Ub4 > M1-Ub4 > HOTag6-G10-Ub, indicating that HOTag6-G10-Ub is preferentially recruited to the dense phase and K48-Ub4 is the least likely to be in the dense phase. Consistently, the Ub4 dense phase concentrations followed the opposite trend of K48-Ub4 < K63-Ub4 < M1-Ub4 < HOTag6-G10-Ub. Interestingly, like for UBQLN2, the Ub dense phase concentrations did not vary with changing Ub:UBQLN2 ratios. However, to our surprise, the Ub:UBQLN2 ratio in the dense phase is much lower than 1:1. Even for HOTag6-G10-Ub, which has the highest dense phase concentration out of the four Ub4, the Ub:UBQLN2 ratio is only about 0.4:1. The substoichiometric Ub:UBQLN2 ratio inside the dense phase is maintained to at least 4:1 total Ub:UBQLN2 ratio (Fig. 5D). These observations indicate that, at a given point, more than half of the UBQLN2 UBA sites are still unbound, i.e. free from ligand binding, in the dense phase. We surmise that Ub-unbound UBA stickers are still needed for interactions with other UBQLN2 regions to drive LLPS. UBQLN2 is a scaffold that undergoes LLPS via homotypic interactions. Upon heterotypic interactions with highly accessible and flexible multivalent ligands via its UBA stickers, UBQLN2 LLPS can be enhanced (Fig. 7A). Hence, the interplay between homotypic and heterotypic interactions is essential for how multivalent ligands that bind to sticker sites on scaffold can modulate LLPS.

**Figure 6.**
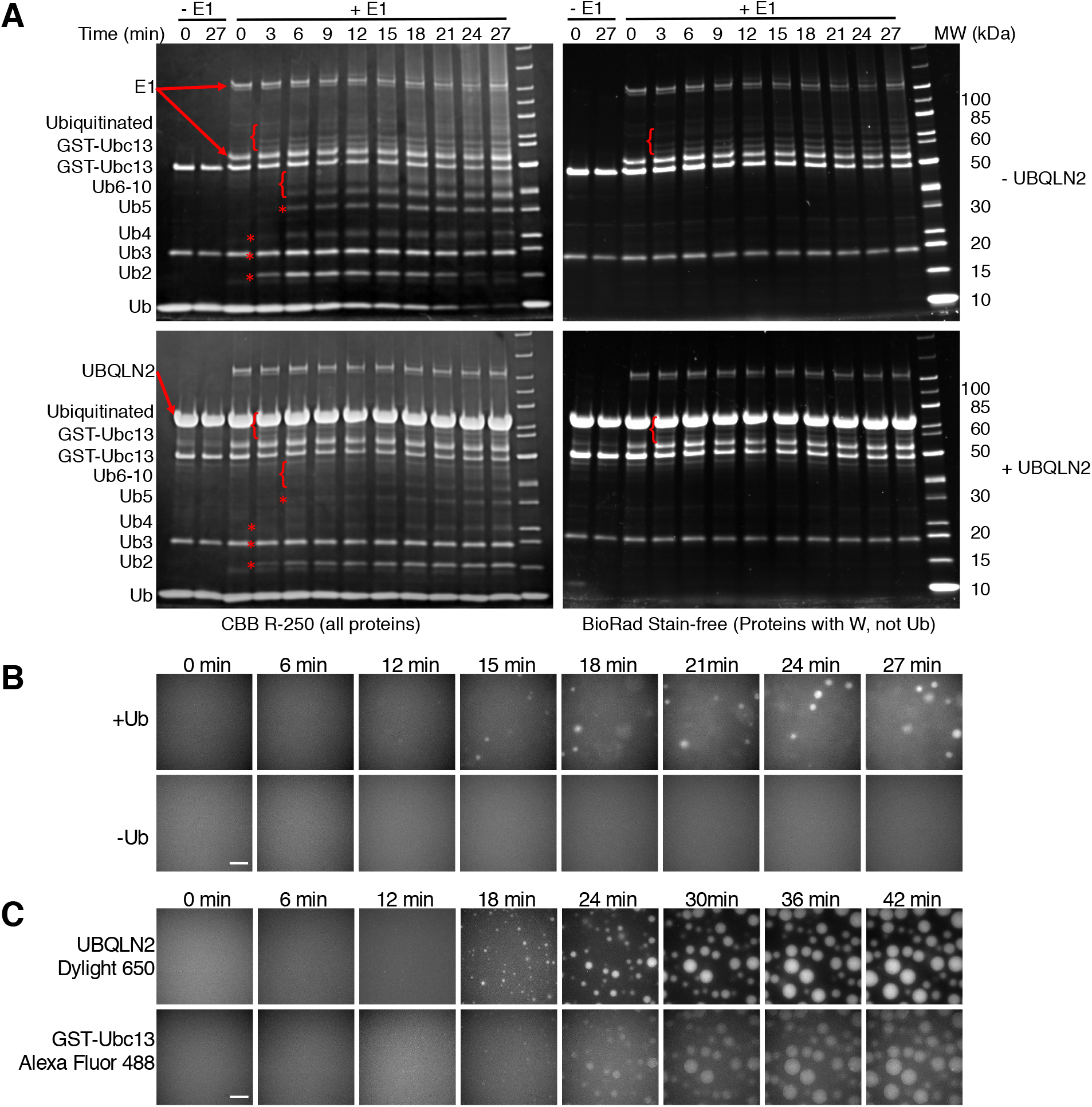
LLPS of UBQLN2 is induced during *in vitro* K63-selective ubiquitination assays. (A) SDS-PAGE gels monitoring the formation of K63-linked polyUb chains and possibly ubiquitinated GST-Ubc13 over time, as indicated by asterisks and curly brackets. K63-Ub3 migrates on the gel similarly as His-Mms2, right below 20 kDa. Experimental conditions: ± 50 μM Ub and UBQLN2, 30 nM Dylight 650-labeled UBQLN2, 1 μM mE1, 2 μM His-Mms2, 4 μM GST-Ubc13, 10 mM ATP and MgCl_2_, 3 mM TCEP, 50 mM Tris pH8, 37 °C. The K63-linked polyUb chain reactions were done in the absence (top) and presence (bottom) of UBQLN2. Since UBQLN2 competes with E1 and E2 to bind to Ub, the presence of UBQLN2 slowed down chain formation (indicated by comparing disappearance rates of Ub). The band intensity for GST-Ubc13 decreased with time. The time-dependent appearance of species between 50-85 kDa were obstructed in the presence of UBQLN2. These species are most likely ubiquitinated GST-Ubc13, and not free polyUb chains, since these bands were observed with BioRad Stain-free gels (right) that detect proteins with tryptophan residues unlike Ub or polyUb chains. (B) Time-lapse fluorescence microscopy monitoring UBQLN2 droplet formation in the presence of K63-selective ubiquitination machinery and with or without Ub. Experimental conditions were the same as in (A). Imaging was done in solution above the coverslip. Scale bar, 5 μm. (C) Time-lapse fluorescence microscopy monitoring formation of UBQLN2- and GST-Ubc13-containing droplets in the presence of Ub and K63-selective ubiquitination machinery. Experimental conditions were the same as in (A) with addition of 30 nM Alexa Fluor 488-labeled GST-Ubc13. Imaging was done at the coverslip surface. Scale bar, 5 μm.

**Figure 7.**
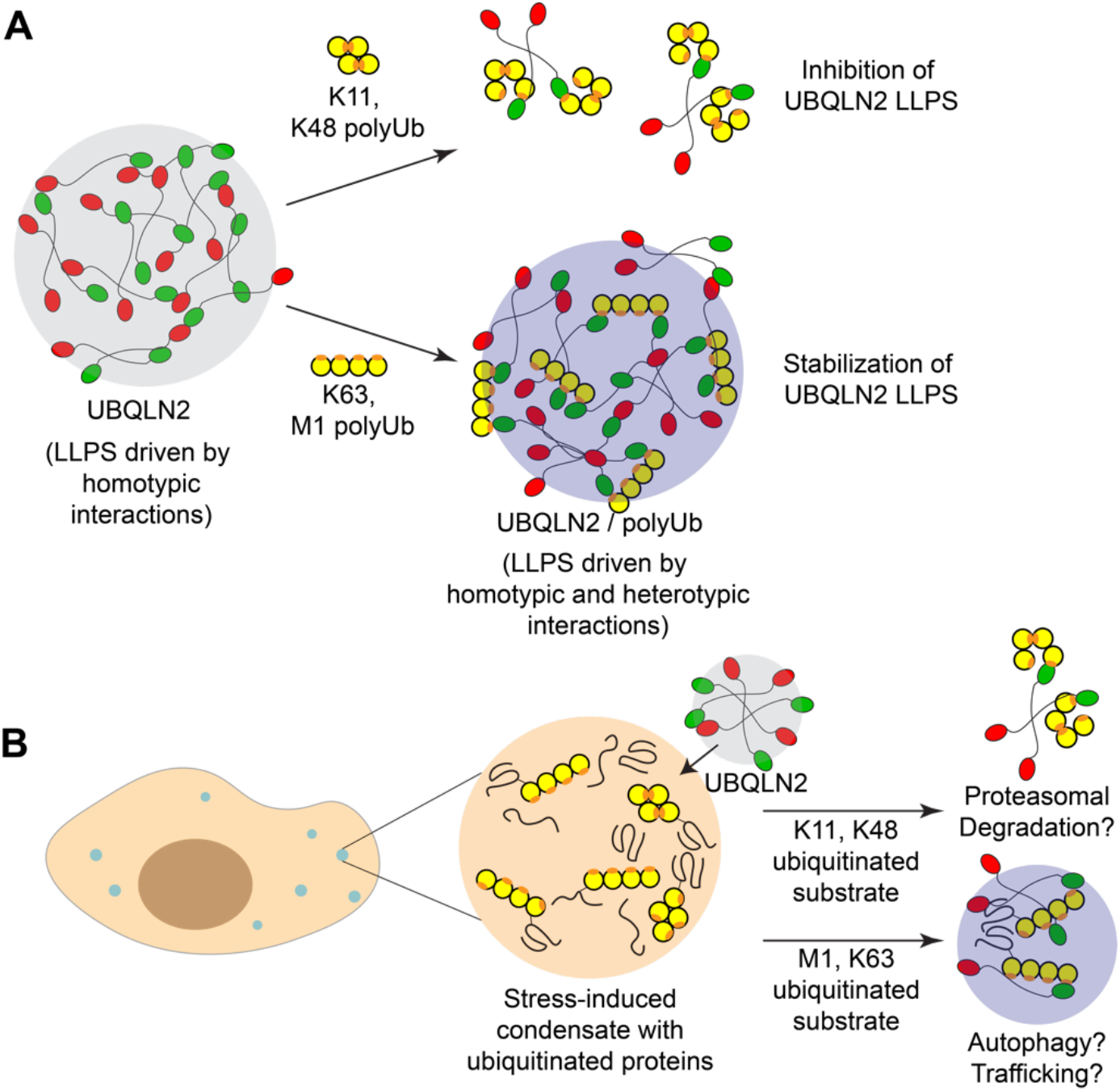
PolyUb chains either inhibit or stabilize LLPS via heterotypic interactions. (A) Compact polyUb chains (e.g. K11, K48) generally inhibit UBQLN2 LLPS, whereas extended chains with accessible hydrophobic patches promote heterotypic LLPS. Orange hotspots on Ub units indicate hydrophobic patch. (B) Model for how polyUb-modulated effects on LLPS may drive different cellular outcomes and signal different PQC pathways for ubiquitinated proteins in stress-induced condensates. K11 and K48 chains typically signal for proteasomal degradation, whereas extended K63 and M1 chains are implicated in autophagy and other non-proteolytic pathways.

The differential partitioning of UBQLN2 and Ub4 into dilute and dense phases can also be observed by fluorescence microscopy (Fig. 5E). UBQLN2 localized mostly inside the droplets at both Ub:UBQLN2 ratios (0.1:1 and 1:1) for all four chains. However, at low Ub:UBQLN2 ratio (0.1:1), K63-Ub4, M1-Ub4 and HOTag6-G10-Ub were also mainly observed inside the droplets, whereas K48-Ub4 was only slightly enriched in the droplets compared to outside. In contrast, at higher Ub:UBQLN2 ratio (1:1), K63-Ub4, M1-Ub4, and HOTag6-G10-Ub were enriched inside the droplets, whereas K48-Ub4 was similarly distributed in the dense and dilute phases. These qualitative observations are entirely consistent with our measured values for the dilute and dense phase concentrations whereby K63-Ub4, M1-Ub4 and HOTag6-G10-Ub preferentially bind UBQLN2 in the dense phase and are recruited to UBQLN2 droplets. These three chains also slowed down the diffusivity dynamics of UBQLN2 in UBQLN2/chain condensates, as indicated by the increased fluorescence recovery after photobleaching (FRAP) half-times (Fig. S8). The slowed dynamics likely resulted from these chains forming an interacting network with UBQLN2, preventing UBQLN2 from moving as freely inside the droplet. However, the three chains had little effects on droplet material properties, as indicated by the similar mobile fractions. K48-Ub4, which has little or no preference to be in UBQLN2 droplets, did not affect the dynamics or material properties of UBQLN2 droplets.

### UBQLN2 condensates assemble during in vitro enzymatic assembly of K63-linked chains

Our data indicate that Ub and polyUb with different lengths and linkages can inhibit or enhance LLPS to different extents. However, in the cell, multiple Ub species, free and conjugated to substrate proteins, are in dynamic equilibrium with one another, depending on the cell cycle, cellular environment and external stimuli (Park and Ryu, 2014). To test how changes in the distribution of different Ub species affect UBQLN2 LLPS, we devised a simplified system comprising 50 μM each of purified Ub and UBQLN2 (spiked with Dylight 650-labeled UBQLN2) as well as K63-specific ubiquitination machinery, including Ub-activating enzyme E1, and Ub-conjugating enzymes Mms2 and GST-Ubc13 (yeast version of UBE2V2-UBE2N). In the presence of MgCl_2_, ATP, TCEP at 37 °C, the amount of free monoUb decreased slowly over time while K63-Ub2, Ub3, Ub4, Ub5 chains were assembled as visible on the gel around 6, 9, 12 and 15 minutes, respectively (Fig. 6A, lower left). Notably, after 27 minutes, free monoUb is still the predominant Ub species, as could be an equilibrium state for many cell types (Kaiser et al., 2011). To complement the monitoring of Ub chain formation by gel, we also observed the mixture under a microscope. At early time points, the solution was homogeneous but after about 15 minutes, micron-sized droplets appeared and grew bigger in size and number over time (Fig. 6B, Movie S1). As a negative control, no droplets were observed in that time frame without Ub in solution (Fig. 6B) or in a solution with Ub and K48-specific ubiquitination machinery (Fig. S9). Therefore, even in the presence of a high ratio of free monoUb to K63-linked polyUb, UBQLN2 can still undergo LLPS as the ratio of different longer Ub chains increases.

Aside from the formation of free K63-linked polyUb chains, we also observed possible formation of ubiquitinated GST-Ubc13 as the intensity of free GST-Ubc13 reduced over time (Fig. 6A). Ub-binding proteins, like Ubc13, can be ubiquitinated in the presence of only E1 and E2 (Hoeller et al., 2007). There seemed to be several species of ubiquitinated GST-Ubc13, most likely mono-ubiquitinated at multiple lysine residues. We wondered if GST-Ubc13 could enter UBQLN2 condensates that form in the presence of K63-selective ubiquitination machinery. To test this, we set up the same experiment as above, spiked it with a small amount of Alexa Fluor 488-labeled GST-Ubc13 and observed droplet formation over time (Fig. 6C). Excitingly, we could see slight enrichment of GST-Ubc13 in UBQLN2 droplets. Most GST-Ubc13 was not ubiquitinated and therefore not localized to UBQLN2 droplets. Ubiquitinated GST-Ubc13 seen here might act like a ubiquitinated substrate that can enter UBQLN2 droplets or even enhance UBQLN2 LLPS before being shuttled to the next step in a PQC pathway. Together, these data indicate that a change in the equilibrium of Ub species (free, conjugated monoUb and polyUb chains) in response to a certain cellular signal can affect UBQLN2 LLPS ability, and perhaps functions in PQC.

## Discussion

Recent studies have highlighted the important role of Ub and polyUb chains in modulating or driving LLPS of Ub-binding shuttle proteins hHR23B, UBQLN2 and p62 (Dao and Castañeda, 2020; Dao et al., 2018; Sun et al., 2018; Yasuda et al., 2020; Zaffagnini et al., 2018). However, these results appear to be at odds with each other. For example, K48-Ub4 mostly disassembles UBQLN2 droplets but drives hHR23B LLPS. Long K63 chains drive p62 LLPS but are much less effective at driving hHR23B LLPS compared to K48 chains. Our mechanistic understanding of how five types of polyUb chains interact with UBQLN2 reconcile these differences and, together with other studies, offer general principles governing the driving force of polyUb chains on shuttle protein LLPS.

We showed that K48-linked chains of four or fewer Ub units generally inhibited or barely enhanced UBQLN2 LLPS whereas K63-linked chains enhanced LLPS over a wide range of Ub:UBQLN2 ratios. We postulated that differences in either chain conformations or binding affinities among UBQLN2 and these chains lead to these different behaviors. We demonstrated, using additional K11, M1-Ub4 chains and HOTag6-G10-Ub, that more extended chains with easily accessible Ub hydrophobic patches can readily promote UBQLN2 LLPS. K63- and M1-linked chains also stabilize LLPS with other proteins, such as p62 (Sun et al., 2018; Zaffagnini et al., 2018). Even though K_d_ values for UBQLN2 and K48-Ub4 differed significantly, depending on the methods employed and whether the labelled partner was UBQLN2 or K48-Ub4, we believe, together with results for K11-Ub4, that these differences stemmed from the compact conformation of K48-Ub4, which has to open for UBQLN2 binding. Moreover, although the binding affinities between UBQLN2 and HOTag6-G10-Ub, M1-Ub4, and K63-Ub4 were similar (Fig. 4G), HOTag6-G10-Ub greatly reduced c_sat_ for UBQLN2 compared to K63-Ub4. Therefore, our data suggested that binding affinities do not contribute significantly to the differences in the effects of polyUb chains on UBQLN2 LLPS.

Interestingly, pulldowns of K48- and K63-linked chains using GST-tagged UBQLN1, a close UBQLN2 homolog that also undergoes LLPS (Gerson et al., 2021), show that FL UBQLN1 only interacts with longer K63-linked chains (Harman and Monteiro, 2019). However, UBQLN1 and UBQLN2 UBAs have no significant preference for either chain (Fig. 4G) (Raasi et al., 2005; Zhang et al., 2008). We speculate that K63-, but not K48-linked, chains undergo LLPS with UBQLN1 during the pulldown process and thus appears to be a tighter binding partner. Like UBQLN2, p62 preferentially phase separates with K63- and M1-linked chains (Sun et al., 2018; Zaffagnini et al., 2018). Also like UBQLN2, FL p62 can only bind to K63-linked chains through pulldown assays (Cabe et al., 2018), whereas p62 UBA bind to both chains with similar affinities (Long et al., 2008; Raasi et al., 2005). Could the preferential pulldown of K63-linked chains by p62 also be an artifact of LLPS? Direct determination of binding affinities and mechanisms between p62 and these chains is needed to tease out the contributions of chain conformation and binding affinity to differences in how p62 phase separates in the presence of different chains. Unlike UBQLN2 and p62, hHR23B appears to bind more tightly to and phase separates preferentially with K48-linked chains than K63-linked chains (Nathan et al., 2013; Raasi et al., 2005; Yasuda et al., 2020). We do note that affinities between hHR23B and polyUb chains need to be confirmed via multiple techniques in light of our conflicting NMR and anisotropy results. Phase diagrams under multiple conditions are needed to conclusively state that K48-linked chains are more efficient at driving hHR23B LLPS. Together, these data indicate that when binding affinities are similar, extended chains with easily accessible Ub hydrophobic patches can stabilize LLPS of Ub-binding shuttle proteins more efficiently. However, preferential binding between a shuttle protein and a compact chain might compensate for the chain’s less extended structure and promote LLPS more efficiently. Moreover, our “designed Ub4”, HOTag6-G_10_-Ub with a long linker between HOTag6 and individual Ub units, resembles a multi-monoubiquitinated substrate. Therefore, substrates that are post-translationally modified with multi-monoubiquitination could drive UBQLN2 LLPS and potentially other Ub-binding adaptor proteins.

Through promoting or inhibiting LLPS, polyUb chains could be potent modulators of signaling outcomes in the cell, akin to PARylation, another PTM that regulates LLPS via length and branching (Reber and Mangerich, 2021). For example, both K11- and K48-linked chains are linked to proteasomal degradation pathways, with a branched K11/K48 chain binding more tightly to the proteasome (Boughton et al., 2019). Intriguingly, only these chain types destabilize UBQLN2 LLPS. In contrast, K63- and M1-linked polyUb are typically involved in non-proteolytic signaling pathways such as cargo sorting, DNA repair, NF-Kβ signaling, and autophagy (Haglund and Dikic, 2012; Linares et al., 2013; Olzmann et al., 2007; Piper et al., 2014; Sun et al., 2018). UBQLN2 has known affiliations with the proteasome, autophagy receptors (LC3) and cargo sorting adaptor proteins (epsin 1 and 2) (Lin et al., 2021; Zheng et al., 2020). We showed that UBQLN2 LLPS can be rapidly modulated by changes in cellular signaling (e.g. formation of K63-linked chains or ubiquitinated substrates, Fig. 6). The UBQLN2 concentration in cells could be below the threshold needed for LLPS such that UBQLN2 is diffuse under normal conditions. In response a cellular event, such as stress, substrates can be ubiquitinated and elicit different responses from UBQLN2, depending on the types of ubiquitination (Fig. 7). UBQLN2 can bind to K11- or K48-linked ubiquitinated substrates, stay diffuse, and shuttle substrates to the proteasome for degradation. UBQLN2 can bind to K63- or M1-linked substrates, colocalize to or form substrate-containing condensates. These condensates can be similar to p62- or hHR23B-containing nuclear degradation foci to which proteasomes are recruited (Fu et al., 2021; Yasuda et al., 2020) or initiation centers for the formation of autophagosomes (Sun et al., 2018; Turco et al., 2019; Zaffagnini et al., 2018). The polyUb effects on UBQLN2 LLPS may be the general mechanism regulating the roles of UBQLN2 and other Ub-binding shuttle proteins in the cell.

UBQLN2 is also recruited to stress granules, most likely due to its LLPS properties (Alexander et al., 2018; Dao et al., 2018). UBQLN2 expression levels are negatively correlated to the number and sizes of stress granules (Alexander et al., 2018), indicating a role for UBQLN2 in regulating the disassembly of stress granules. Stress granules are cleared either following the removal of K63-linked polyubiquitinated G3BP1, a stress granule core protein, or by autophagy, depending on the stressor types and length of exposure (Buchan et al., 2013; Gwon et al., 2021). Through differential modulation of UBQLN2 LLPS by different polyUb chains, UBQLN2 may facilitate these different stress granule clearance processes (Fig. 7). Persistent stress granules can lead to the formation of disease-linked protein-containing inclusions (Nedelsky and Taylor, 2019), that comprise, among others, PQC components such as UBQLN2 and polyUb chains (Ceballos-Diaz et al., 2015; Deng et al., 2011; Sharkey et al., 2020). Therefore, knowledge of how polyUb chains affect UBQLN2 in stress granules is essential to understanding the cellular functions of UBQLN2.

Aside from cellular implications, our work presented here also contributes to the important question of how binding to multivalent ligands modulates the phase boundaries of phase-separating systems. Using monovalent and divalent ligands, Ruff and colleagues demonstrated through simulations that high-valency ligands that bind more tightly to spacer sites (residues or protein regions not involved in driving LLPS) are more effective at lowering the c_sat_ necessary for LLPS of multivalent scaffolds (Ruff et al., 2021a). Conversely, ligands that bind to sticker sites generally increase c_sat_, but the effect is lessened with increasing ligand valency. Our experimental work recapitulates many of these observations and offers additional insights. We systematically and quantitatively demonstrated that longer polyUb could either less effectively inhibit (K48 chains) or more effectively promote (K63 chains) UBQLN2 LLPS (Fig. 1). In fact, K48-Ub4 slightly promoted UBQLN2 LLPS at low Ub:UBQLN2 ratios. Following this trend, K48 chains with five or more Ub units would further enhance UBQLN2 LLPS, in similar manner as K63 chains with 3 or 4 Ub units. K48 and K63 polyUb ligands interact mainly with UBQLN2 UBA stickers, but K48-linked chains also interact with an additional sticker site. Therefore, K48 chains need to have increased valency (by increasing chain length) to compensate for the loss of a sticker site important for UBQLN2 self-interactions and LLPS. We expect multivalent ligands that bind to sticker sites of other systems to enhance LLPS when a certain ligand valency is reached.

Despite having the same valency and binding to the same UBQLN2 stickers (Fig. 4C) with similar affinities (Fig. 4H), the ligands K63-Ub4, M1-Ub4 and HOTag6-G10-Ub lower c_sat_ for UBQLN2 LLPS to highly different extents (Fig. 5D). The major difference among these chains is how the UBQLN2 binding surface is presented (i.e. the spacer length and flexibility between the stickers). Therefore, how a multivalent ligand modulates LLPS of a scaffold (e.g. UBQLN2) depends not only on binding affinities, ligand valencies, and binding sites on scaffold, but also on the properties of the spacers between the stickers on the ligand. In our case, it appears that longer, more flexible spacers are better at promoting LLPS, likely due to the ligand generating a broader interacting network. However, there must be a limit after which the multivalent ligand begins to behave like individual units of monovalent ligands. It would be interesting to determine what that limit is in terms of spacer length / flexibility and if this principle can be generalized to other systems.

Overall, we uncovered the molecular mechanisms enabling distinct polyUb chains to be potent regulators of phase separation of UBQLN2 and possibly other Ub-binding shuttle proteins. With this knowledge, we can design specific experiments in cells, such as inhibition of certain polyUb chains or PQC pathways, to elicit how polyUb chains may tune UBQLN2’s ability to form or be recruited into condensates and then carry out PQC functions. Moreover, disease mutations in UBQLN2 have been shown to disrupt UBQLN2’s roles in PQC (Chen et al., 2018; Halloran et al., 2020; Hjerpe et al., 2016). Therefore, determining the effects of polyUb chains on the phase separation of UBQLN2 variants is an essential next step to elucidating disease-related mechanisms involving UBQLN2.

## Acknowledgements

This work was supported by ALS Association grant 18-IIP-400, NIH R01 GM136946 (turbidity assays, microscopy, and NMR experiments) and NSF CAREER MCB 1750462 (SEC-MALS-SAXS experiments) to C.A.C. NMR data were acquired on an 800 MHz NMR spectrometer funded by NIH shared instrumentation grant 1S10OD012254. FRAP data were acquired at the Syracuse University Blatt BioImaging Center on a Zeiss LSM980 with Airyscan2 funded by NIH S10 OD026946-01A1. This research used resources of the Advanced Photon Source, a U.S. Department of Energy (DOE) Office of Science User Facility operated for the DOE Office of Science by Argonne National Laboratory under Contract No. DE-AC02-06CH11357. This project was supported by grant P30 GM138395 from the National Institute of General Medical Sciences of the National Institutes of Health. Use of the Pilatus 3 1M detector was provided by grant 1S10OD018090-01 from NIGMS. We thank Chris Waudby for insightful conversations on binding affinities, and Rohit Pappu, Kiersten Ruff, Tanja Mittag, Daniel Kraut, Elliot Dine, and Susan Krueger for stimulating discussions over the years leading to this project. We also thank Ashley Canning with assistance on AUC experiments. The content is solely the responsibility of the authors and does not necessarily reflect the official views of the National Institutes of Health.

## Author Contributions

Conceptualization, T.P.D. and C.A.C.; Methodology, T.P.D., Y.Y., J.H., S.N.L, C.A.C.; Investigation, T.P.D., Y.Y., M.S.C., W.M., M.F.P.; Writing – Original Draft, T.P.D., C.A.C.; Writing – Review & Editing, T.P.D., M.S.C., S.N.L, C.A.C; Funding Acquisition, C.A.C.; Resources, C.A.C., M.S.C., J.H., W.M., S.N.L; Supervision, C.A.C.

## Methods

### Gene synthesis and site-directed mutagenesis

The HOTag6-G10-Ub polypeptide has the following sequence: MTLREIEELLRKIIEDSVRSVA ELEDIEKWLKKIGGGGGGGGGGMQIFVKTLTGKTITLEVEPSDTIENVKAKIQDKEGIPPDQQRLIFAGKQLEDGRTLSDYNIQKESTLHLVLRLRGG. The genes encoding HOTag6-G10-Ub and M1-Ub4-His6 were codon-optimized, synthesized and cloned into pET24b (Novagen) by GenScript (NJ, USA). K48R, K63R, K11/63R Ub and Ub-V-His6 were made using Phusion Site-Directed Mutagenesis Kit (Thermo Scientific).

### Protein Expression, and Purification

Wildtype, K48R, K63R, Ub and Ub-V-His6 were expressed and purified as detailed elsewhere (Beal et al., 1996; Castañeda et al., 2016b). The gene encoding mouse E1 was a kind gift from Jorge Eduardo Azevedo (Addgene plasmid 32534, (Carvalho et al., 2011)). E1, Mms2, Yuh1 and GST-Ubc13 were expressed in *Escherichia coli* NiCo21 (DE3) cells in Luria-Bertani (LB) broth at 16 °C overnight. GST-E2-25K in pGEX-4T2 and GST-Ube2s-UBD in pGEX6P1 were expressed in Escherichia coli Rosetta 2 (DE3) pLysS cells in Luria-Bertani (LB) broth at 16 °C overnight. Bacteria were pelleted, frozen, then lysed via freeze/thaw method in 50 mM Tris, 1 mM EDTA (pH 8), 1 mM PMSF, 1 mM MgCl_2_, and 25 U of Pierce universal nuclease. Yuh1, E1 and Mms2 were purified via Ni^2+^ affinity chromatography. GST-E2-25K, GST-Ubc13, and GST-Ube2s-UBD were purified via GST chromatography. All proteins were concentrated, buffer exchanged into 50 mM Tris and 1 mM EDTA (pH 8), and stored at 80 °C for subsequent use in the production of K48- and K63-linked polyUb chains, and K11-linked Ub4.

K48-linked and K63-linked Ub2, Ub3 and Ub4 were synthesized sequentially. Briefly, equal amounts of K48R (K63R) Ub and Ub-V-His6 incubated with 1000 nM E1 and 10 μM GST-E2-25K (2 μM His-Mms2 and 4 μM GST-Ubc13) in the presence of 10 mM ATP, 0.3 mM TCEP in Tris buffer at pH 8 for 3 hours at 37 °C. This procedure generates K48R (K63R) Ub2 with the C-terminal end of the proximal Ub blocked by V-His6. Yuh1 was added to remove the V-His6 from the end of Ub2, which was then purified via cation exchange column using 50 mM ammonium acetate (pH 4.5) as the buffer. Protein was eluted via a linear gradient from 0 to 100% of 50 mM ammonium acetate, 1 M NaCl (pH 4.5). Purified Ub2 was then buffer exchanged into 50 mM Tris buffer at pH 8. K48 and K63-Ub3 were made the same way Ub2 was made. K48 and K63-Ub4 required an additional purification step via size exclusion chromatography using a Superdex 75 HiLoad 16/600 column (GE Healthcare). K11-Ub2 was made with K63R as described by (Bremm et al., 2010) using mE1 and GST-Ube2s-UBD in similar conditions as for K48 and K63 chains. K11-Ub4 was made from K11-Ub2 and purified by size exclusion chromatography using a Superdex 75 HiLoad 16/600 column (GE Healthcare). The yield for K11-Ub4 was about 10%, much lower than for K48 and K63-Ub4, which was about 50%.

HOTag6-G10-Ub and M1-Ub4-His6 were expressed in *Escherichia coli* NiCo21 (DE3) cells in Luria-Bertani (LB) broth at 37 °C overnight. Bacteria were pelleted, frozen, then lysed via freeze/thaw method in 50 mM Tris, 1 mM EDTA (pH 8), 1 mM PMSF, 1 mM MgCl_2_, and 25 U of Pierce universal nuclease. The cleared lysate for HOTag6-G10-Ub was then loaded onto anion exchange column and eluted with a gradient between 20 mM HEPES, 0.02% NaN_3_ (pH 7) and 20 mM HEPES (pH 7), 1 M NaCl, 0.02% NaN_3_. Fractions containing HOTag6-G10-Ub were dialyzed, loaded onto a cation exchange column, and eluted with a gradient between 50 mM ammonium acetate (pH 4.5) and 50 mM ammonium acetate, 1 M NaCl (pH 4.5). Purified HOTag6-G10-Ub was then concentrated, and buffer exchanged into 20 mM NaPhosphate, 0.5 mM EDTA, 0.1 mM TCEP, 0.02% NaN_3_ (pH 6.8). M1-Ub4-His6 was purified via Ni^2+^ chromatography, dialyzed into 50 mM Tris, 1 mM DTT before the addition of Yuh1 to cleave the His-tag after the last Ub unit. The cleaved M1-Ub4 was then dialyzed into 50 mM ammonium acetate pH 4.5, loaded onto cation exchange column, and eluted with a gradient between 50 mM ammonium acetate (pH 4.5) and 50 mM ammonium acetate, 1 M NaCl (pH 4.5). The fractions containing M1-Ub4 were concentrated and exchanged into 20 mM NaPhosphate, 0.5 mM EDTA, 0.1 mM TCEP, 0.02% NaN_3_ (pH 6.8). Purified proteins were frozen at −80°C.

Full-length UBQLN2 and UBQLN2 450-624 were expressed and purified as described by previously (Dao et al., 2018). Briefly, the constructs were expressed in *E. coli* Rosetta 2 (DE3) pLysS cells in Luria-Bertani (LB) broth at 37°C overnight. Bacteria were pelleted, frozen, lysed, then purified via a “salting out” process. NaCl was added to the cleared lysate to the final concentration of 0.5 M - 1 M. UBQLN2 droplets were pelleted and then resuspended in 20 mM NaPhosphate, 0.5 mM EDTA, 0.1 mM TCEP, 0.02% NaN_3_ (pH 6.8). Leftover NaCl was removed through HiTrap desalting column (GE Healthcare). Purified proteins were frozen at −80°C.

### Fluorescent Labeling

For imaging, determination of dense and dilute phase concentrations and FRAP experiments, UBQLN2, GST-Ubc13, K48-Ub4, K63-Ub4, M1-Ub4, HOTag6-G10-Ub constructs were fluorescently labeled with Dylight-488, DyLight-650 (Thermo Scientific) or Alexa Fluor-488 NHS Ester (Molecular Probes), according to the manufacturer’s instructions. For fluorescence anisotropy experiments, UBQLN2 450-624 with a S624C mutation, K48-Ub4 and K63-Ub4 with an S20C mutation on the most distal Ub were fluorescently labeled with Alexa Fluor 488 C5 Maleimide (Invitrogen). Excess dye was removed via size exclusion chromatography using an ENrich™ SEC 650 10 x 300 Column (BioRad).

### Spectrophotometric Absorbance/Turbidity Measurements

Protein samples were prepared by mixing protein solution containing UBQLN2 and polyUb (the concentrations of the protein stock solutions were doubled compared with the sample concentrations) and cold sodium phosphate buffer (pH 6.8, 20 mM NaPhosphate, 0.5 mM EDTA, 0.1 mM TCEP, 0.02% NaN_3_) containing 400 mM NaCl at 1:1 stoichiometry. The protein concentrations were chosen to cover as wide a range as possible to allow observation of phase separation during the temperature ramps. All the solutions were kept on ice for at least 5 minutes before mixing. Absorbance at 600 nm was recorded as a function of temperature by a Cary 3500 UV-Vis spectrophotometer using a temperature ramp rate of 1 °C/min increasing from 16 °C to 60 °C (4°C to 60 °C range was screened in some cases). Net absorbance values were recorded after subtracting the absorbance value of a buffer control. At least two trials (n = 2) were conducted to ensure reproducibility. Data were plotted using Mathematica (Wolfram Research).

### Phase Diagram Measurements

For the LCST (lower critical solution temperature) phase transition, i.e. mapping the phase boundary as temperature is increased, protein samples were prepared as described for the turbidity measurements. Cloud point temperatures were determined by fitting a Four Parameter Logistic Regression model to the data:

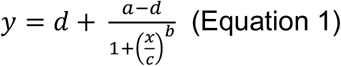

Cloud point temperatures (T_cp_) used were the points of inflection (c). Cloud point temperatures were used to define the coexistence curve as a function of ligand:protein ratio (ligand refers to polyUb chains and protein refers to UBQLN2). Fitting and plotting of data were done with Mathematica (Wolfram Research) and Kaleidagraph (Synergy Software).

### Bright-field Imaging of Phase Separation

Samples were prepared to contain 50 μM UBQLN2 and different amounts of Ub and Ub chains in 20 mM NaPhosphate, 200 mM NaCl, 0.1 mM TCEP, and 0.5 mM EDTA (pH 6.8), spiked with UBQLN2 labeled with Dylight 650 and/or Ub chains labeled with Dylight 488. Samples were added to Eisco Labs Microscope Slides, with Single Concavity, and covered with MatTek coverslips that had been coated with 5% bovine serum albumin (BSA) to minimize changes due to surface interactions, and incubated coverslip-side down at 30 °C for 10 min. Phase separation was imaged on an ONI Nanoimager (Oxford Nanoimaging Ltd, Oxford, UK) equipped with a Hamamatsu sCMOS ORCA flash 4.0 V3 camera using an Olympus 100×/1.4 N.A. objective. Images were prepared using Fiji (Schindelin et al., 2012) and FigureJ plugin.

### Calculations of Dilute and Dense Phase Concentrations

Samples were prepared on ice to contain 50 μL of 70 μM UBQLN2, 10 μM of UBQLN2 labeled with Alexa Fluor 488, 5 μM of Ub or Ub chains labeled with Dylight 650, and different amounts of Ub and Ub chains in 20 mM NaPhosphate, 200 mM NaCl, 0.1 mM TCEP, and 0.5 mM EDTA (pH 6.8). Samples were incubated at 30 °C for 10 minutes, then centrifuged at 10000 x g for 5 minutes at 30 °C. Without disrupting the pellet, as much of the supernatant as possible was transferred to a new tube. 8 μL of 8 M urea solution was added to the pellet. Tube containing pellet and urea was incubated for 1 hour at room temperature, vortexed and centrifuged. The volume of the mixture was determined by pipetting with a P10. The dense phase volume was determined by subtracting the total volume by 8 μL of urea added. Absorbance at 493 nm (for Alexa Fluor 488/UBQLN2) and 655 nm (for Dylight 650/Ub chains) were recorded. Dense and dilute phase concentrations were determined with a standard curve for multiple concentrations of UBQLN2 mixed with UBQLN2 labeled with Alexa Fluor 488 and Ub chains mixed with Ub chains labeled with Dylight 650.

### Fluorescence Recovery After Photobleaching (FRAP)

To perform FRAP on UBQLN2/Ub chain droplets of similar size, samples were prepared to contain different UBQLN2 concentrations, depending on Ub chain types and molar ratios, in 20 mM NaPhosphate, 200 mM NaCl, 0.1 mM TCEP, and 0.5 mM EDTA (pH 6.8). Specifically, for UBQLN2-only K48-Ub4/UBQLN2 at 1:1 samples, UBQLN2 concentration is 60 μM. For K48-Ub4/UBQLN2 at 1:1, UBQLN2 concentration is 75 μM. For K63-Ub4/UBQLN2 at 0.5:1 and 1:1, UBQLN2 concentration is 50 μM. For K63-Ub4/UBQLN2 at 2:1, UBQLN2 concentration is 60 μM. For M1-Ub4/UBQLN2 at 0.5:1 and 1:1, UBQLN2 concentration is 40 μM. For M1 Ub4/UBQLN2 at 2:1, UBQLN2 concentration is 50 μM. For HOTag6-Ub/UBQLN2 at 0.5:1 and 1:1 and 2:1, UBQLN2 concentration is 30 μM. Samples were added to Eisco Labs Microscope Slides, with Single Concavity, and covered with MatTek coverslips that had been coated with 5% bovine serum albumin (BSA), and incubated coverslip-side down at 30 °C for 20-30 min. FRAP was carried on a Zeiss LSM 980 with Airyscan 2 confocal microscope (Carl Zeiss AG, Oberkochen, Germany) using a Plan-Apochromat 63X/1.4 NA oil. Images were prepared using Fiji (Schindelin et al., 2012) and FigureJ plugin.

### NMR Spectroscopy

Proteins were prepared in 20 mM NaPhosphate buffer (pH 6.8), 0.5 mM EDTA, 0.02% NaN_3_, and 5% D_2_O. ^1^H-^15^N SOFAST-HMQC was used for titration experiments of UBQLN2 constructs and polyUb, although additional ^1^H-^15^N TROSY-HSQC experiments were collected at distinct Ub:UBQLN2 ratios. ^1^H-^15^N TROSY-HSQC experiments were used for experiments involving fulllength UBQLN2. All NMR experiments were performed on a Bruker Avance III 800 MHz spectrometer equipped with a TCI cryoprobe at 25°C. ^1^H-^15^N SOFAST-HMQC experiments were acquired using spectral widths of 16 and 30 ppm in the direct ^1^H and indirect ^15^N dimensions, using 2048 and 160 total points, respectively. NMRPipe (Delaglio et al., 1995) was used to process all NMR data, and peak analyses were performed using CCPNMR 2.5 (Vranken et al., 2005). Chemical shift perturbations (CSPs) were quantified as follows:

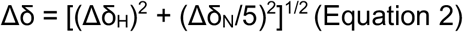

where Δδ_H_ and Δδ_N_ are the differences in ^1^H and ^15^N chemical shifts, respectively.

### NMR Titration Experiments and K_d_ Determination

Unlabeled protein ligand (Ub4 in most cases) was titrated into 50 μM samples of ^15^N-labeled protein (UBQLN2), and the binding was monitored by recording ^1^H-^15^N TROSY-HSQC or SOFAST-HMQC spectra as a function of ligand:protein ratios of concentrations. Concentrations of Ub ligand were adjusted by the number of Ub units in the polyUb chain such that 50 μM Ub4 was equal to 200 μM Ub. For TITAN (Waudby et al., 2016) lineshape analysis, only SOFAST-HMQC spectra for titrations involving ^15^N-labeled protein (UBQLN2 450-624) were used, following the NMR processing protocol in (Waudby and Christodoulou, 2020). For all cases, the single-site binding model (two-state ligand binding in TITAN) was used. Parameters were first estimated using a K_d_ of 10 μM and a k_off_ rate of 5000 s^-1^. Binding affinity calculations were performed using global fitting of peaks from a total of 10-12 amide resonances (Supplementary Figure S6). We evaluated how well the peaks were fit using two-dimensional contour maps, and removed poorly fit peaks (typically in crowded areas of the spectrum). Once complete, errors were evaluated using jackknife analysis; representative best fit of spectra are shown in Supplementary Figure S6.

### Sedimentation Velocity Analytical Ultracentrifugation

Purified UBQLN2 487-624, K48-Ub4, K63-Ub4, or UBQLN2:Ub4 complexes were loaded into 3- or 12-mm two-sector charcoal-filled Epon centerpieces with sapphire windows. All experiments were carried out using a Beckman Coulter ProteomeLab XL-A analytical ultracentrifuge equipped with absorbance optics and a 4-hole An-60 Ti rotor at 60,000 rpm that was pre-equilibrated to 20°C prior to running the experiment. The samples were scanned with a zero second time interval between scans for 300 scans and analyzed by the continuous distribution (*c*(*s*)) method in the program SEDFIT (Schuck, 2000). Concentration profiles (*a*(*r*, *t*)) were modeled as the sum of Lamm Equation solutions scaled by a continuous distribution *c(s)* as follows:

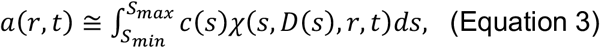

where *s* is the sedimentation coefficient, *χ*(*s*,*D*(*s*),*r*,*t*) is the Lamm Equation solution that is dependent on *D(s)*, the corresponding diffusion coefficient, *r*, radius from the center of rotation, and *t*, the time from the beginning of the experiment (Padrick and Brautigam, 2011; Schuck, 2000). The program SEDNTERP (Laue et al., 1992) was used to calculate the buffer density (1.0064 mL/g), viscosity (0.009 P), and partial specific volume of UBQLN2 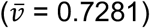, K48-Ub4 (0.746), K63-Ub4 (0.746), K48-Ub4+UBQLN2 (0.7405), and K63-Ub4+UBQLN2 (0.7426), which were based on the amino acid sequences.

Multi-signal sedimentation velocity (MSSV) analysis was performed to determine the stoichiometry of the complex formed between UBQLN2 and Ub4. In MSSV, the standard *c(s)* approach is modified to deconvolute the contributions of individual species with distinct spectral properties in a component distribution *c_k_(s)* where *k* represents the individual components of a mixture. Here, the absorbance at wavelength *λ* (*a_λ_*(*r*, *t*)) is modeled as:

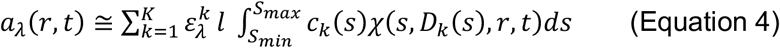

where *l* is the path-length, *K* is the number of solutes present, and *c_k_(s)* is a continuous distribution for component *k*. The spectral properties of the individual proteins were first determined by SV-AUC using absorbance at 260 nm and 280 nm. Global component *c_k_(s)* analysis of both signals was performed using the known *ε^280^* extinction coefficient as a reference to calibrate *ε^260^* for each protein (Table S4). The results indicate good spectral discrimination between UBQLN2 and both Ub4s with fitted spectral discrimination parameter D_*norm*_ > 0.236 (Brautigam et al., 2013). MSSV of the UBQLN2:Ub4 mixtures were then collected at both 260 nm and 280 nm and globally fit using the multiwavelength discrete/continuous distribution analysis with mass constraints in SEDPHAT version 15-2b. Integration of the resulting *c_k_(s)* distributions revealed the molar signal increment for each protein under a peak at a given sedimentation value (Figure S4). All plots were created with GUSSI (Brautigam, 2015).

To determine the statistical significance of the resulting molar ratios, global *c_k_(s)* analyses were conducted by constraining the molar ratio to different values and determining if the resulting fit to the data 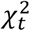 is significantly worse than the unconstrained analysis *χ*^2^, using the built in F-statistics calculator to determine the critical *χ*^2^ at 1σ and 2σ. Fixed molar ratios with 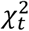 values exceeding the critical *χ*^2^ value at 2σ (dashed red line in Figure S4) were deemed significantly different from that of the unconstrained analysis and rejected (Padrick and Brautigam, 2011).

### Fluorescence polarization anisotropy

Fluorescence anisotropy measurements were made on a SpectraMax i3x plate reader (Molecular Devices) with the FP-FLUO settings. Experiments were done using 100 nM fluor-UBQLN2 450-624 in 20 mM sodium phosphate buffer, 0.5 mM EDTA, pH 6.8 at 25 °C containing 10 μM BSA to reduce protein sticking to the plate. Ub or Ub chains were prepared in the same solution at 150 μM Ub (M1-Ub4, HOTag6-Ub), or 50 μM Ub (K48-Ub4 and K63-Ub4), or 50 μM Ub (K11-Ub4), and 1.5-fold, or 1.5-fold, or 2-fold serial-dilutions, respectively, were prepared in the 96-well plate (100 μL per well, read height 1.18 mm). Reciprocal experiments were done using 100 nM fluor-K48-Ub4 or fluor-K63-Ub4 and 50 μM UBQLN2 450-624 in 2-fold serial-dilutions. A single-site binding equation was used to fit anisotropy data as a function of Ub concentration:

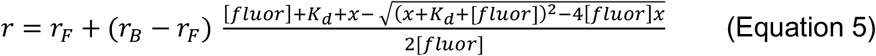

where x is ligand concentration, [*fluor*] is the concentration of fluor-UBQLN2 450-624 or fluor-K48-Ub4 or fluor-K63-Ub4 (100 nM for all titrations), *r* is the observed anisotropy, *r_F_* is the anisotropy at zero ligand concentration and *r_B_* is the anisotropy at saturating ligand concentration. *K*_d_ were determined from an average of three replicates. Fitting and plotting were done with Kaleidagraph (Synergy Software).

### SEC-MALS-SAXS Experiments

SAXS was performed at BioCAT (beamline 18ID at the Advanced Photon Source, Chicago) with in-line size exclusion chromatography (SEC) to separate sample from aggregates and other contaminants thus ensuring optimal sample quality and multiangle light scattering (MALS), dynamic light scattering (DLS) and refractive index measurement (RI)) for additional biophysical characterization (SEC-MALS-SAXS). All protein samples were prepared in 20 mM NaPhosphate buffer at pH 6.8 containing 0.5 mM EDTA and 0.02% NaN_3_ (See Table S4 for details). The samples were loaded on a Superdex 200 10/300 Increase column (Cytiva) at a temperature of 20°C run by a 1260 Infinity II HPLC (Agilent Technologies) at 0.6 ml/min. The flow passed through (in order) the Agilent UV detector, a MALS detector and a DLS detector (DAWN Helios II, Wyatt Technologies), and an RI detector (Optilab T-rEX, Wyatt). The flow then went through the SAXS flow cell. The flow cell consists of a 1.0 mm ID quartz capillary with ~20 μm walls. A coflowing buffer sheath is used to separate sample from the capillary walls, helping prevent radiation damage (Kirby et al., 2016). Scattering intensity was recorded using a Pilatus3 X 1M (Dectris) detector which was placed 3.6 m from the sample giving us access to a q-range of 0.003 Å^-1^ to 0.35 Å^-1^. 0.5 s exposures were acquired every 2 s during elution and data was reduced using BioXTAS RAW 2.1.1 (Hopkins et al., 2017). Buffer blanks were created by averaging regions flanking the elution peak (see Figure S5) and subtracted from exposures selected from the elution peak to create the I(*q*) vs *q* curves used for subsequent analyses. Molecular weights and hydrodynamic radii were calculated from the MALS and DLS data, respectively, using the ASTRA 7 software (Wyatt). Additionally, R_g_ and I(0) values were obtained using the entire q-range of the data by calculating the distance distribution functions, P(r) versus r, using GNOM (Svergun, 1992).

