## Supplementary Material for "Mechanistic insights into the enhancement or inhibition of phase separation by polyubiquitin chains of different lengths or linkages"

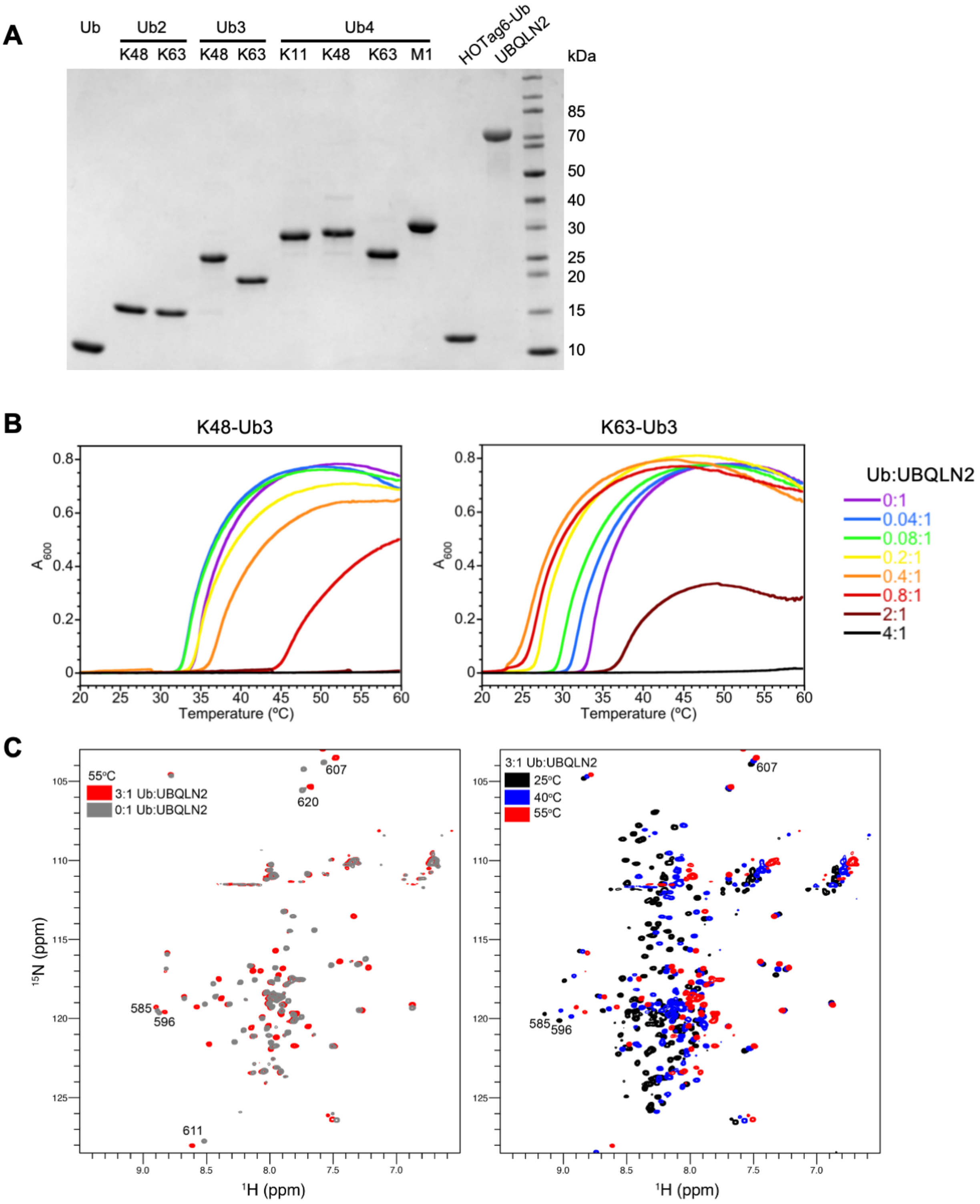


**Supplementary Figure S1**. (A) SDS-PAGE gel of polyUb chains and UBQLN2 used in this study. (B) Representative turbidity traces for solutions of 30 μM UBQLN2 mixed with different amounts of K48-Ub3 or K63-Ub3 polyUb chains. Data show that low amounts of K48-Ub3 (Ub:UBQLN2 ratios < 0.2:1) minimally affect the transition temperature at which $A_{600}$ of a

UBQLN2 solution increases and phase separates, whereas low amounts of K63-Ub3 significantly decrease the temperature threshold at which the UBQLN2 solution phase separates. In contrast, high amounts of both chains increase the temperature for where UBQLN2 solutions phase separate. (C) $^{1}H$-$^{15}N$ CP-HISQC NMR spectra of 50 μM UBQLN2 450-624 with 150 μM Ub collected at 25°C (black), 40°C (blue), and 55°C (red) in pH 6.8 buffer containing 20 mM NaPhosphate with no added NaCl. Spectra at 55°C are compared to spectra of UBQLN2 without any added Ub (gray). Contour settings are identical for each set of spectra. Representative UBA peaks are labeled with corresponding residue number.

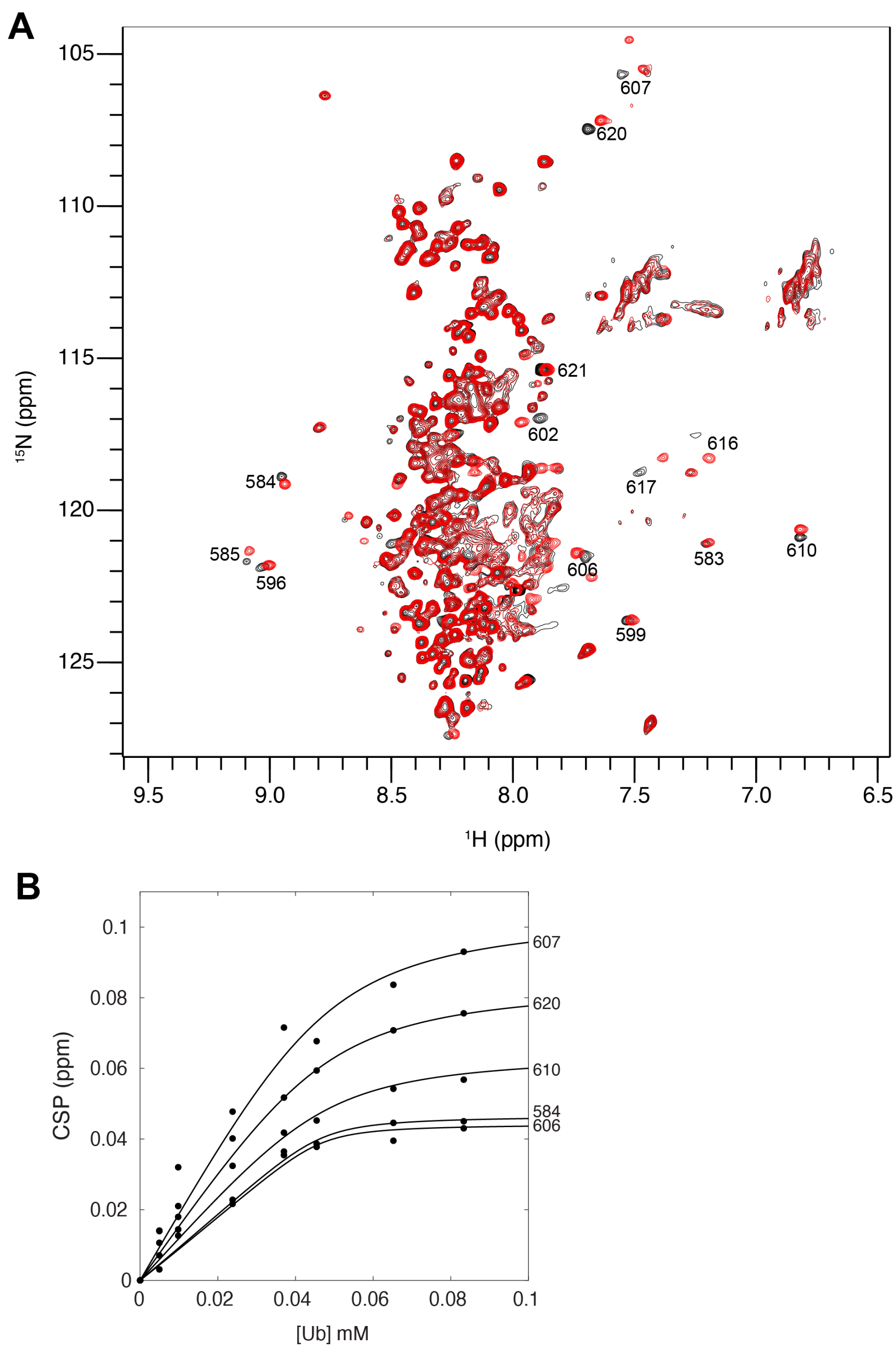


**Supplementary Figure S2.** (A) $^{15}N$-$^{1}H$ TROSY-HSQC spectra of FL UBQLN2 (50 μM) in the absence and presence of Ub (~2:1 Ub:UBQLN2 ratio). Highlighted are those amide resonances in the UBA that exhibited CSPs in the presence of Ub. These same residues are also involved in the self-interactions that are important for UBQLN2 LLPS. Spectra acquired and processed with identical parameters. (B) Residue-specific amide titration curves of UBQLN2. Fits from a single-site binding model were overlaid on experimental data points ($K_d$ ~ 3 μM +/- 2 μM).

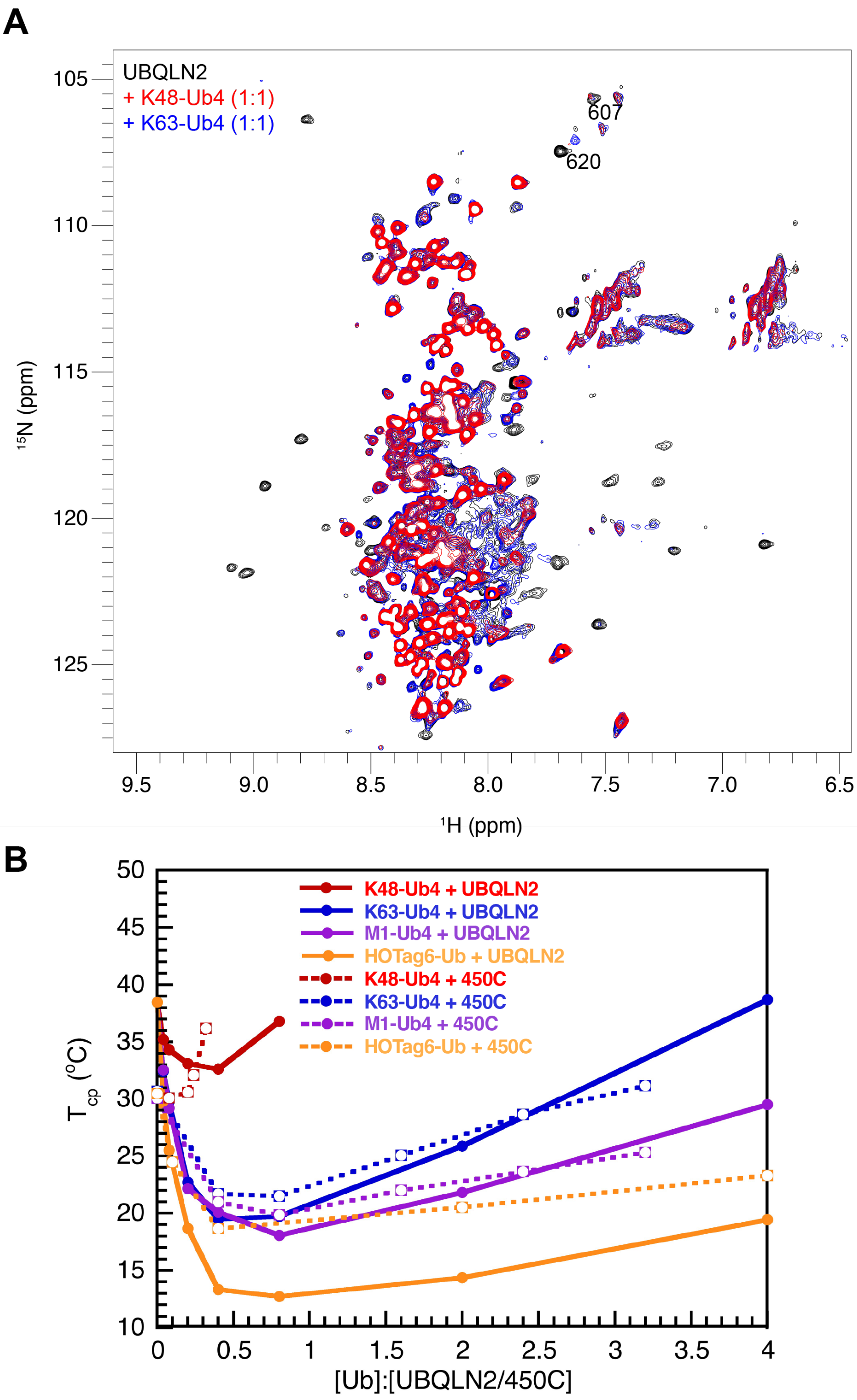


**Supplementary Figure S3.** (A) $^{15}$N-$^{1}$H TROSY-HSQC spectra of FL UBQLN2 (50 μM) in the absence and presence of K48-Ub4 or K63-Ub4 (~1:1 Ub4:UBQLN2 ratio) as noted in the color legend. Note that UBA resonances are no longer visible upon addition of Ub4. Spectra acquired and processed with identical parameters. (B) Temperature-composition phase diagram of FL UBQLN2 (at 30 μM) and UBQLN2 450-624 (at 100 μM) indicating that these two proteins are affected similarly by the addition of four different Ub4 chains. Using solutions containing distinct amounts of polyUb chain mixed with UBQLN2, the temperature midpoint of the phase transitions (cloud point temperatures ($T_{cp}$)) were extracted from spectrophotometric turbidity assays as described in Methods. Solid and dashed line indicates the phase boundary of FL UBQLN2 and UBQLN2 450-624, respectively. Above this line, the solution is phase-separated, and below which the solution is homogeneous.

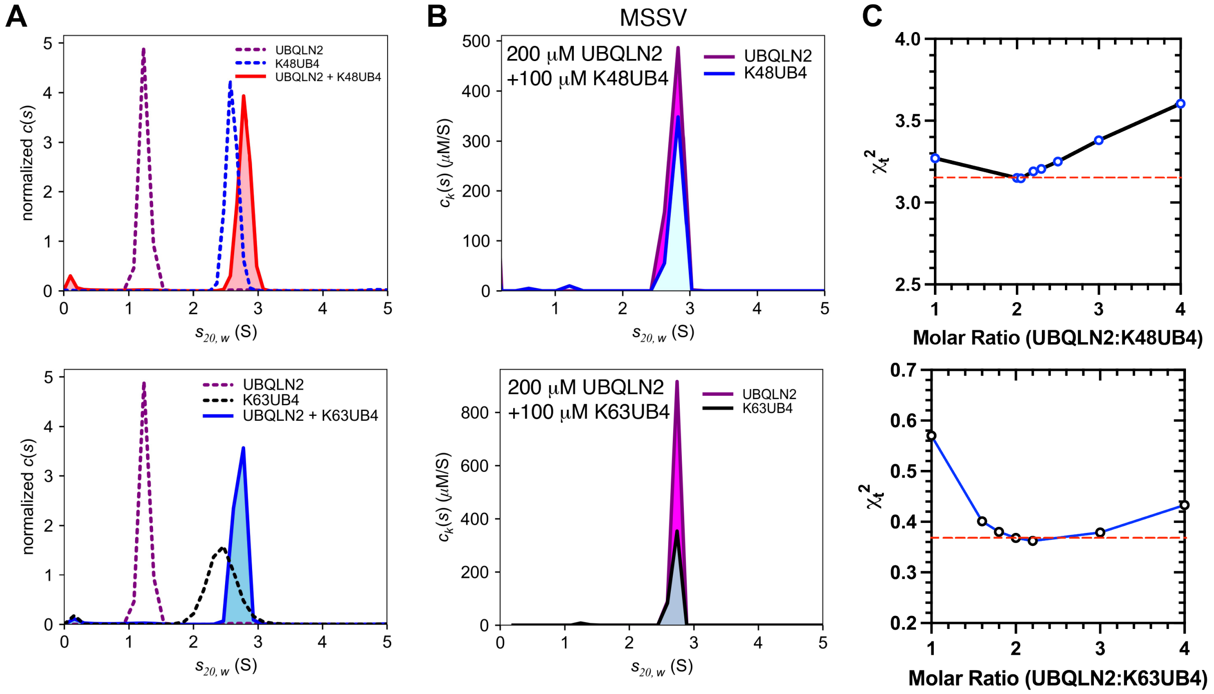


**Supplementary Figure S4**. Sedimentation velocity analytical ultracentrifugation (SV-AUC) experiments of UBQLN2 487-624 in the absence and presence of K48-Ub4 or K63-Ub4. (A) Normalized c(s) distribution for the individual components UBQLN2 487-624 (200 μM) and Ub4 (100 μM), as well as the UBQLN2:Ub4 complex at a loading stoichiometric ratio of 2:1 (this corresponds to a UBQLN2:Ub ratio of 1:2). (B, C) Multi-signal sedimentation velocity (MSSV) analysis to determine stoichiometry of UBQLN2:Ub4 complex with 200 μM UBQLN2 and 100 μM Ub4 (see Tables S3 and S4). Component $c_k(s)$ distributions for UBQLN2 and Ub4 are colored according to legend. No free UBQLN2 is observed in the $c_k(s)$ distribution for either complex. (C) To probe accuracy of calculated stoichiometry of the complex (Table S3), preconstrained values of the molar ratio were tested, and best fit ($\chi_t^2$) values were determined. Experiments show best fit when molar ratio was constrained to an average of two UBQLN2 molecules bound for every Ub4 molecule.

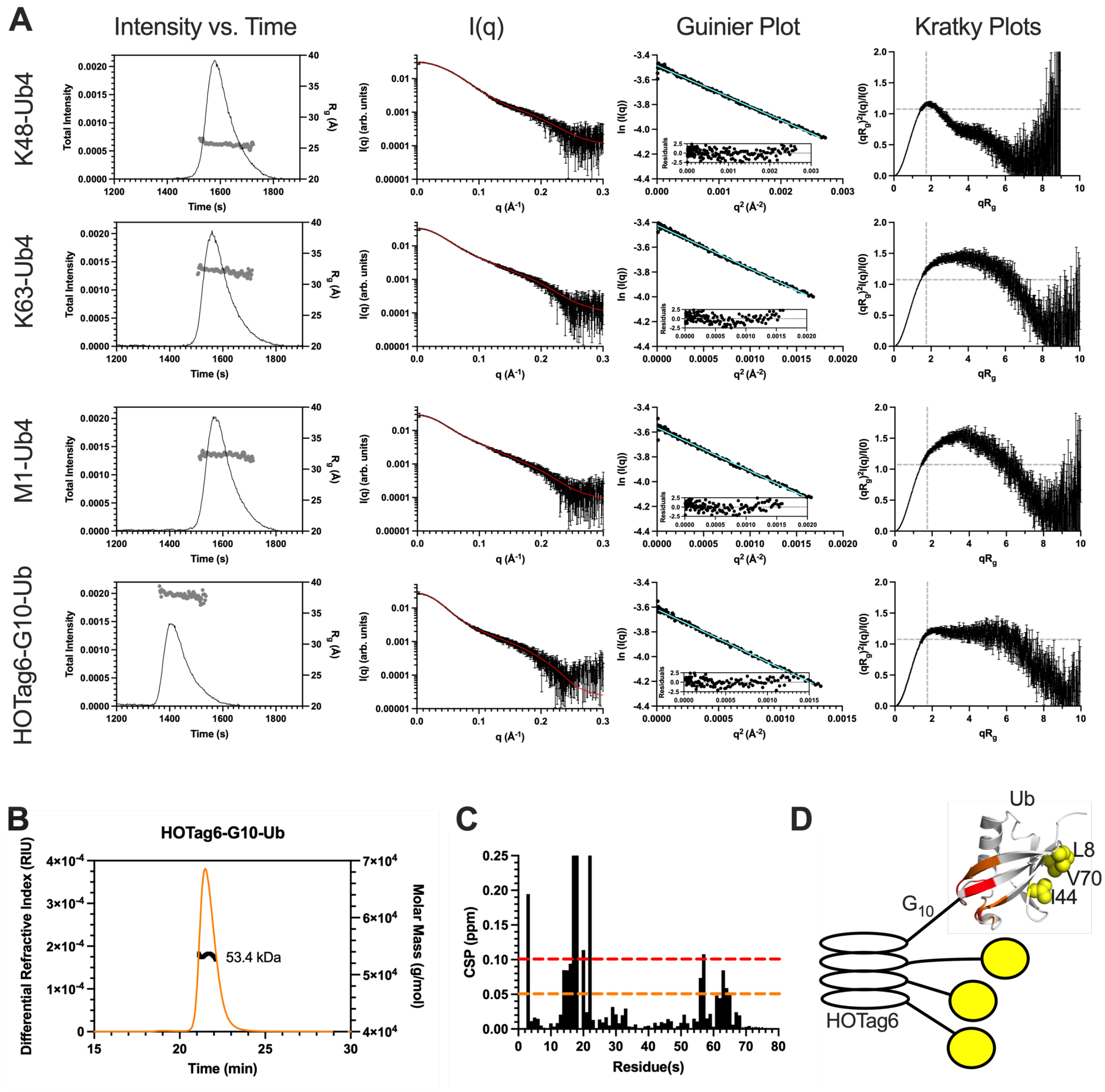


**Supplementary Figure S5.** (A) SEC-SAXS profiles for K48-Ub4, K63-Ub4, M1-Ub4, and HOTag6-G10-Ub. I(q) vs. q scattering curves (middle left) determined from frames 784-808 (K48-Ub4), 766-791 (K63-Ub4), 791-813 (M1-Ub4), and 705-728 (HOTag6-G10-Ub) on the corresponding SEC-SAXS profiles (left). Red line in I(q) profile indicates fit from P(r) analysis (see Fig. 4B). Cyan line in Guinier plot (middle right) is linear fit of ln(I(q)) vs. $q^2$, while inset shows residuals of fit. Dimensionless Kratky plots (right) include dashed lines to indicate where a globular protein would peak. Increase in polyUb chain flexibility from top (K48-Ub4) to bottom (HOTag6-G10-Ub) is indicated by shifts in the peak position up and to the right of the globular peak and larger plateaus in the higher q region. (B) HOTag6-G10-Ub is a tetramer; the molecular weight (MW) of HOTag6-G10-Ub was determined using SEC-MALS-SAXS experiments. The observed MW value of 53.4 kDa is consistent with the expected MW of a HOTag6-G10-Ub tetramer (52.976 kDa), four times that of the monomer (13.244 kDa). (C) CSPs of amide resonances in the Ub unit of HOTag6-G10-Ub vs. monoUb. Note that no significant CSPs exist for the hydrophobic patch region consisting of residues 8, 44, and 70. (D) CSPs from panel B are mapped onto a cartoon model of the HOTag6-G10-Ub tetramer. CSPs are color-coded red (CSPs $\geq$ 0.1 ppm) and orange (CSPs $\geq$ 0.05 ppm). Hydrophobic patch residues 8, 44, and 70 are shown in sphere representation.

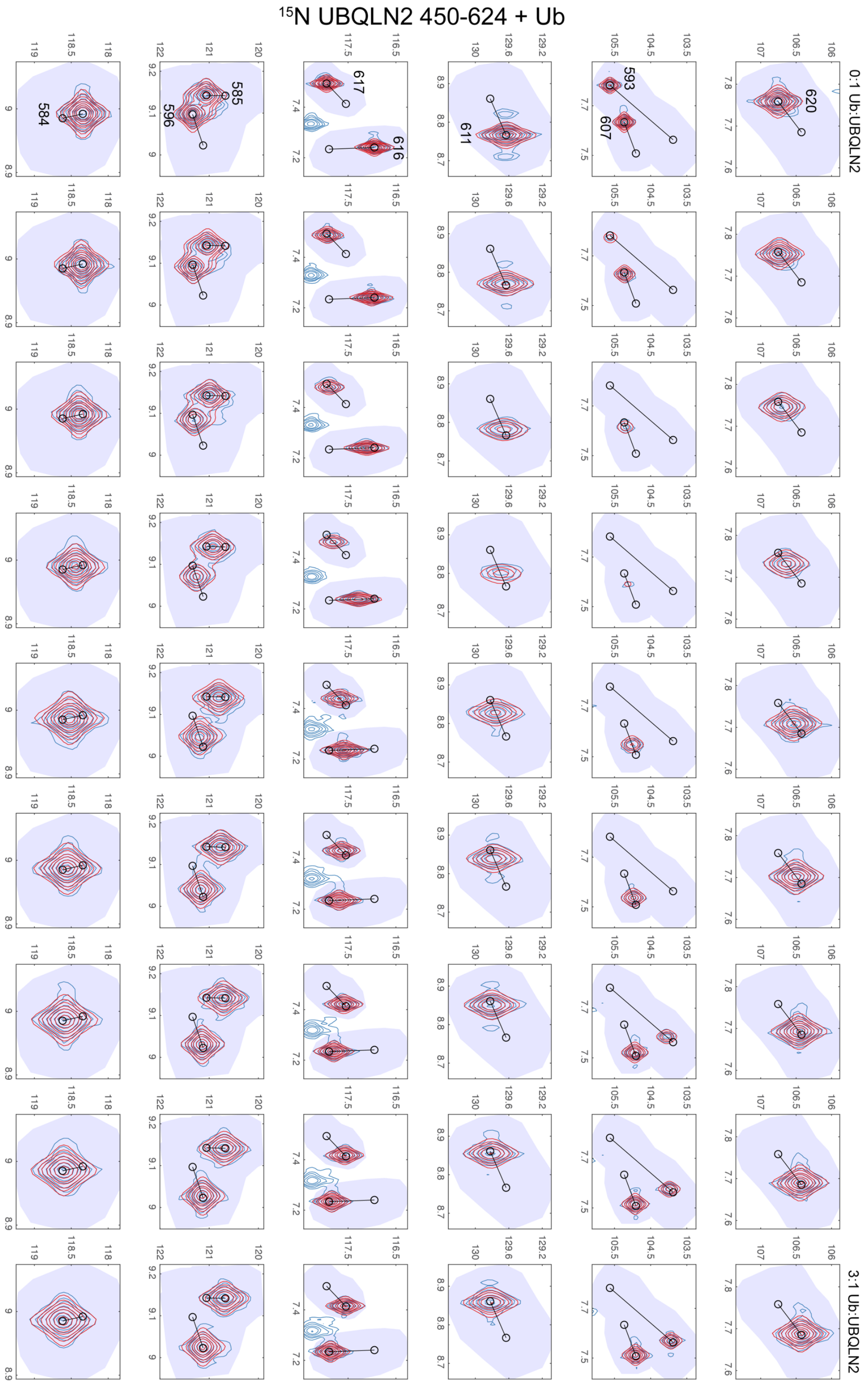


**Supplementary Figure S6A.** TITAN fits using single-site binding equation for UBQLN2 450-624 amide resonance-specific trajectories (residues labeled in 0:1 Ub:UBQLN2 spectra) in the presence of Ub as a function of Ub:UBQLN2 ratio. Original spectra are shown in blue contours, with fitted spectra shown in red contours.

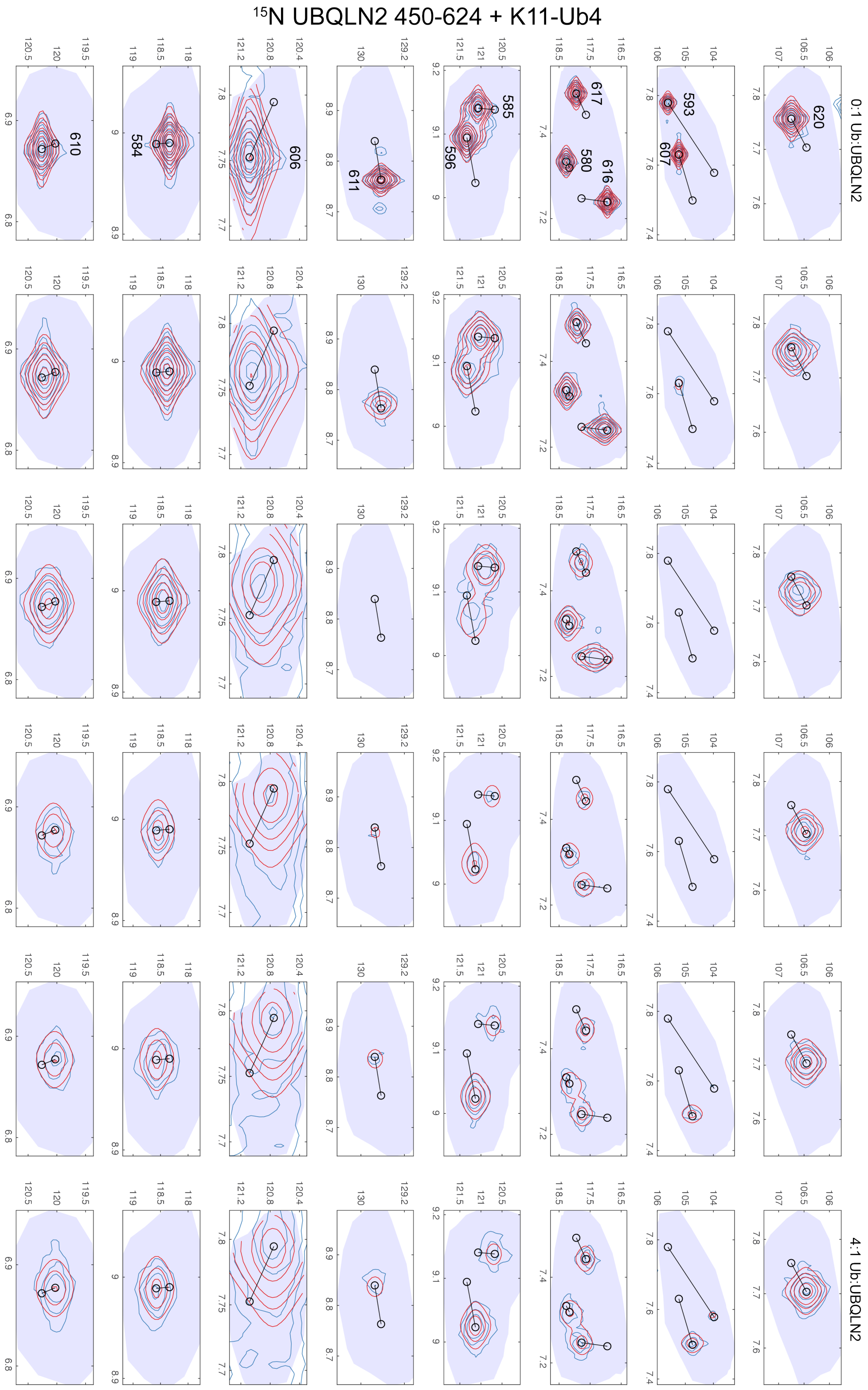


**Supplementary Figure S6B.** TITAN fits using single-site binding equation for UBQLN2 450-624 residue-specific trajectories in the presence of K11-Ub4 as a function of Ub:UBQLN2 ratio.

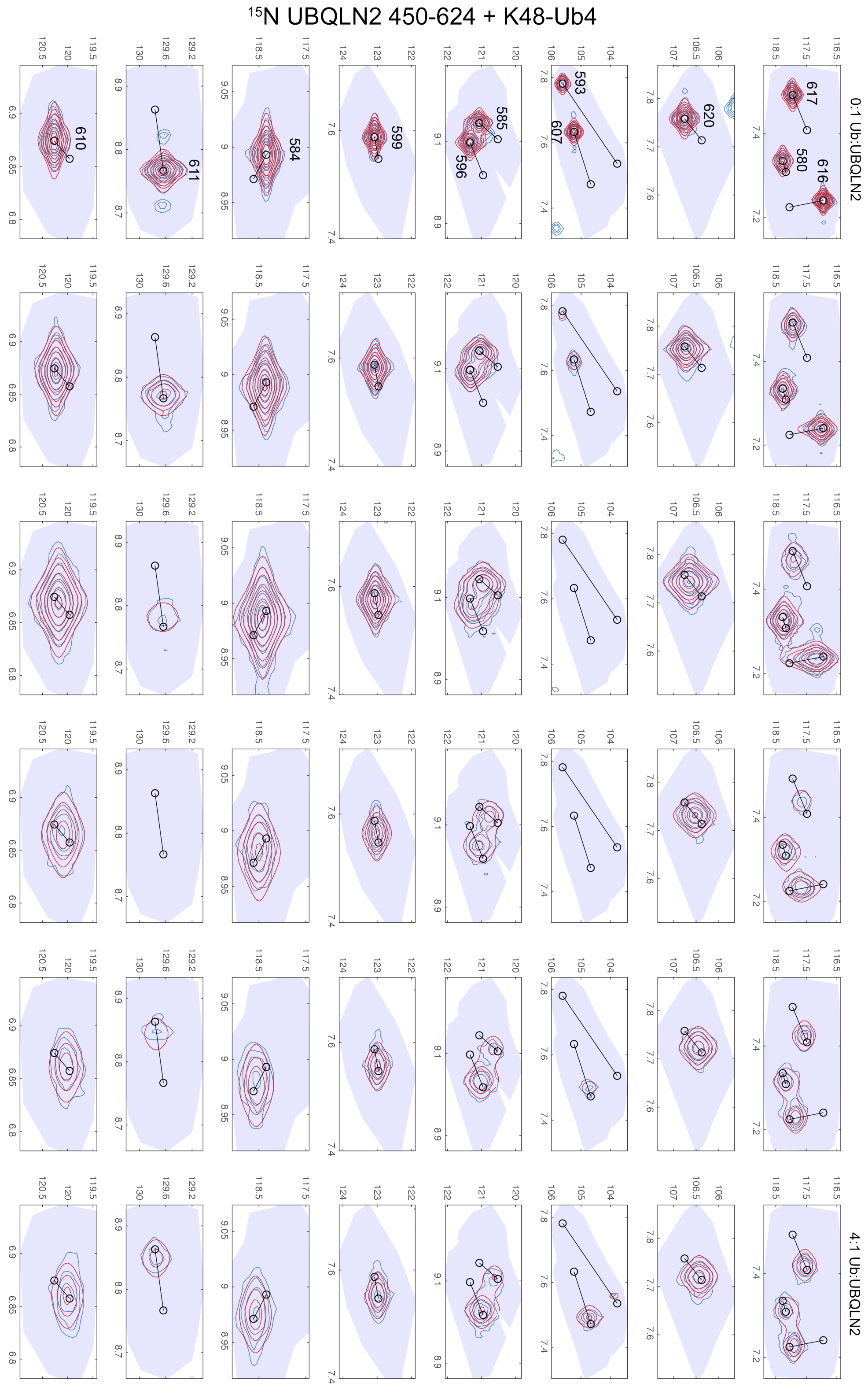


**Supplementary Figure S6C.** TITAN fits using single-site binding equation for UBQLN2 450-624 residue-specific trajectories in the presence of K48-Ub4 as a function of Ub:UBQLN2 ratio.

$^{15}N$ UBQLN2 450-624 + K63-Ub4

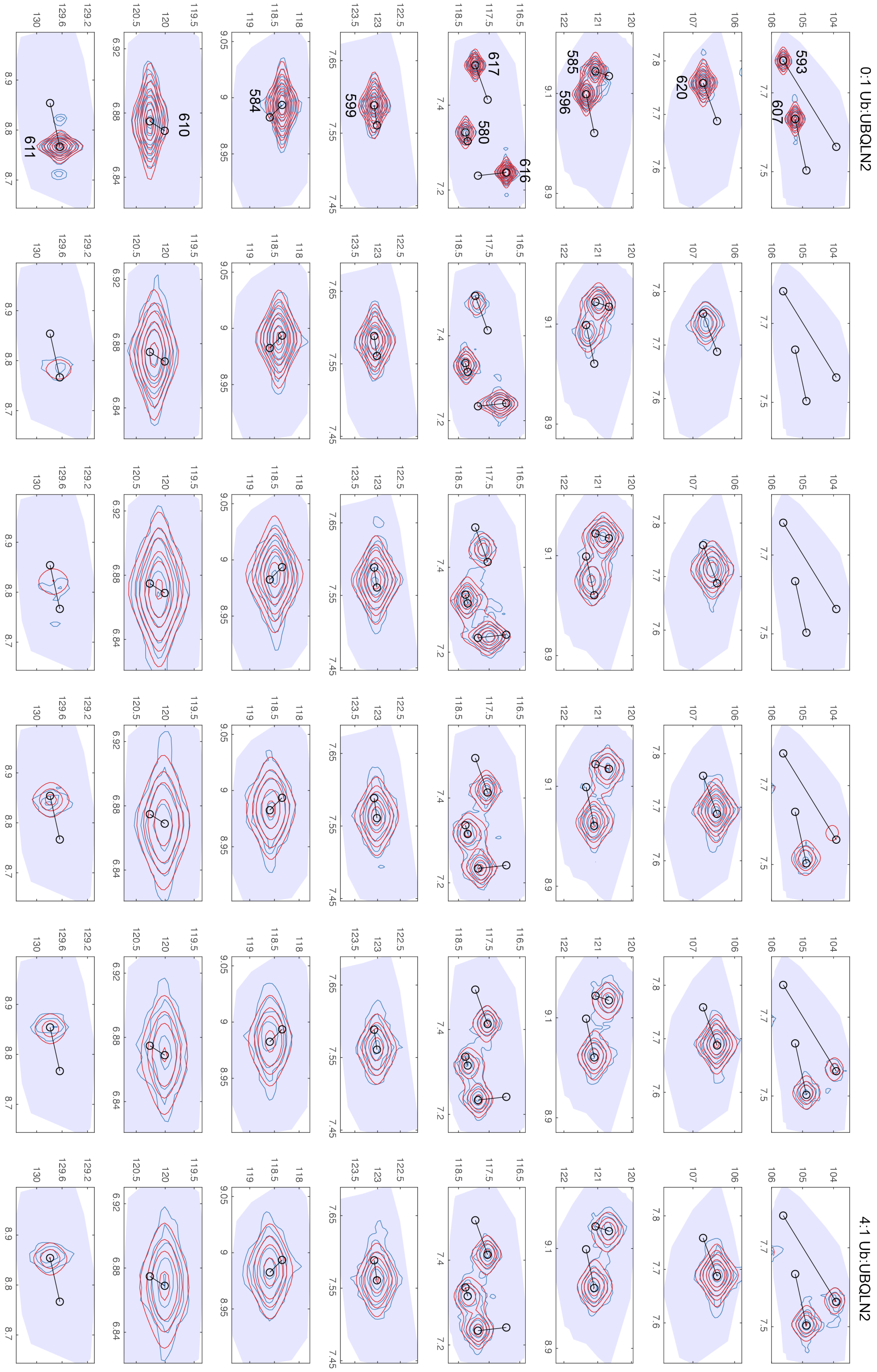


**Supplementary Figure S6D.** TITAN fits using single-site binding equation for UBQLN2 450-624 residue-specific trajectories in the presence of K63-Ub4 as a function of Ub:UBQLN2 ratio.

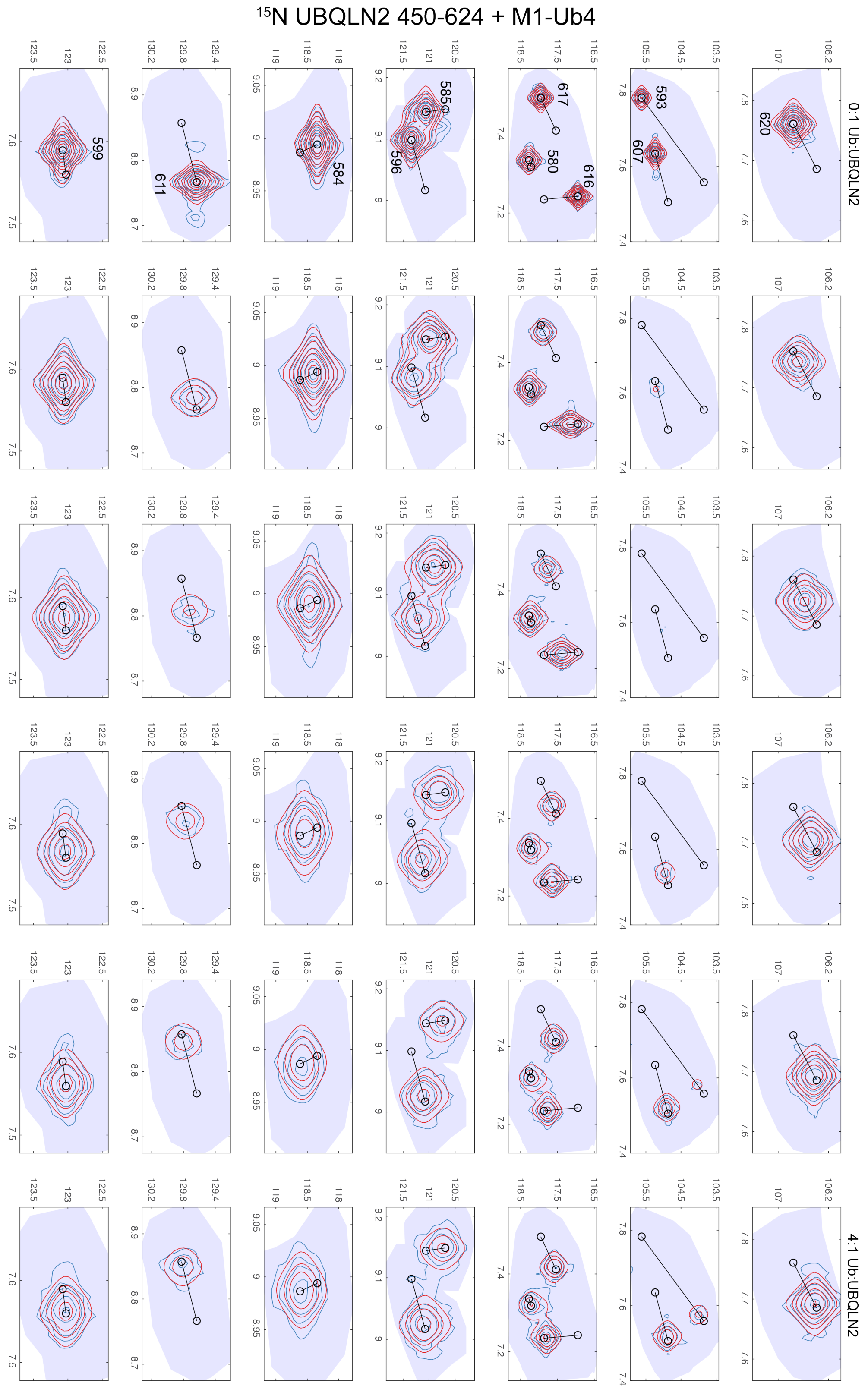


**Supplementary Figure S6E.** TITAN fits using single-site binding equation for UBQLN2 450-624 residue-specific trajectories in the presence of M1-Ub4 as a function of Ub:UBQLN2 ratio.

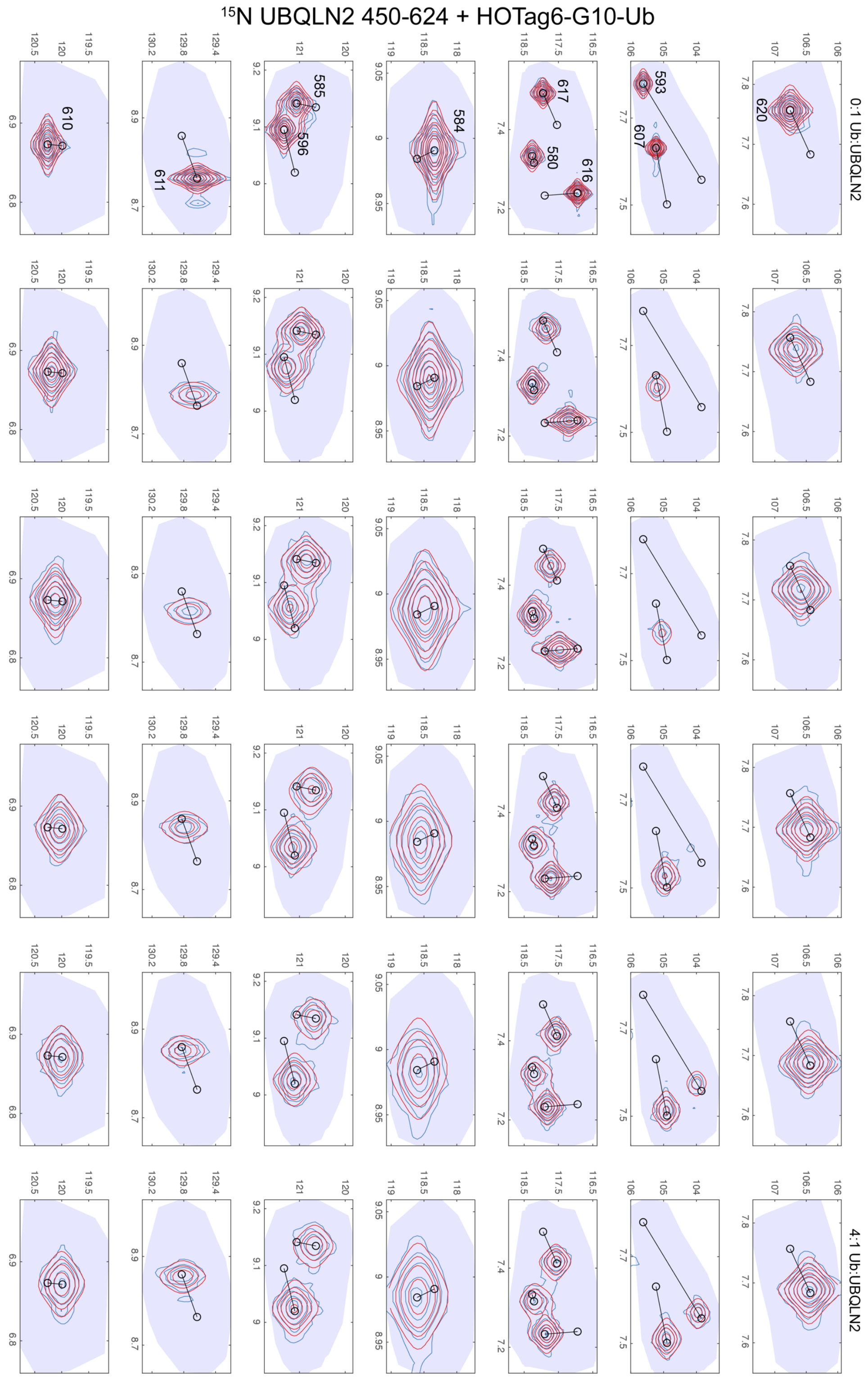


**Supplementary Figure S6F.** TITAN fits using single-site binding equation for UBQLN2 450-624 residue-specific trajectories in the presence of HOTag6-G10-Ub as a function of Ub:UBQLN2 ratio.

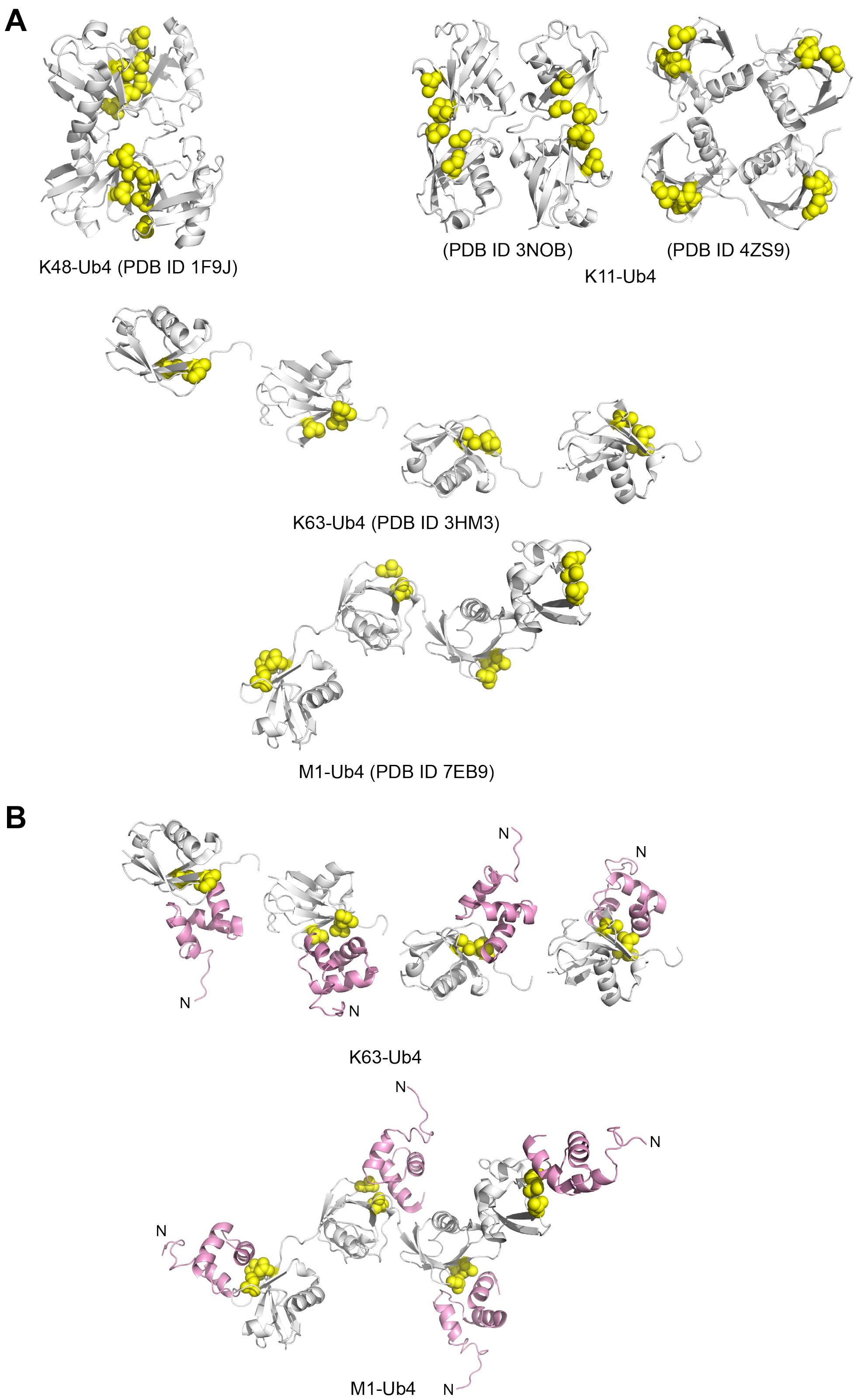


**Supplementary Figure S7.** (A) Representative structures of polyUb tetramers for K48-Ub4, K63-Ub4 and M1-Ub4 (with ligand removed) from PDB entries as noted. Shown are two putative structures of K11-Ub4, based on K11-Ub2 (PDB ID 3NOB) or a model of K11-Ub4 (PDB ID 4Z9S, (Levin-Kravets et al., 2015)) based on K11-Ub2 in PDB ID 2XEW. Shown as yellow spheres are the hydrophobic patch residues L8, I44, and V70 in each Ub unit. Note the compact conformations of both K11-Ub4 and K48-Ub4, but extended conformations for M1-Ub4 and K63-Ub4. (B) Models of how UBQLN2 UBA domains (pink) could dock to either K63-linked or M1-linked polyUb chains. C-terminal UBQLN2 UBA domains were positioned by overlaying the UBQLN1 UBA:Ub complex from PDB ID 2JY6 onto each Ub unit. The UBQLN1 UBA exhibits 97% sequence identity with the UBQLN2 UBA domain. N-termini of each UBA domain are marked with ‘N’.

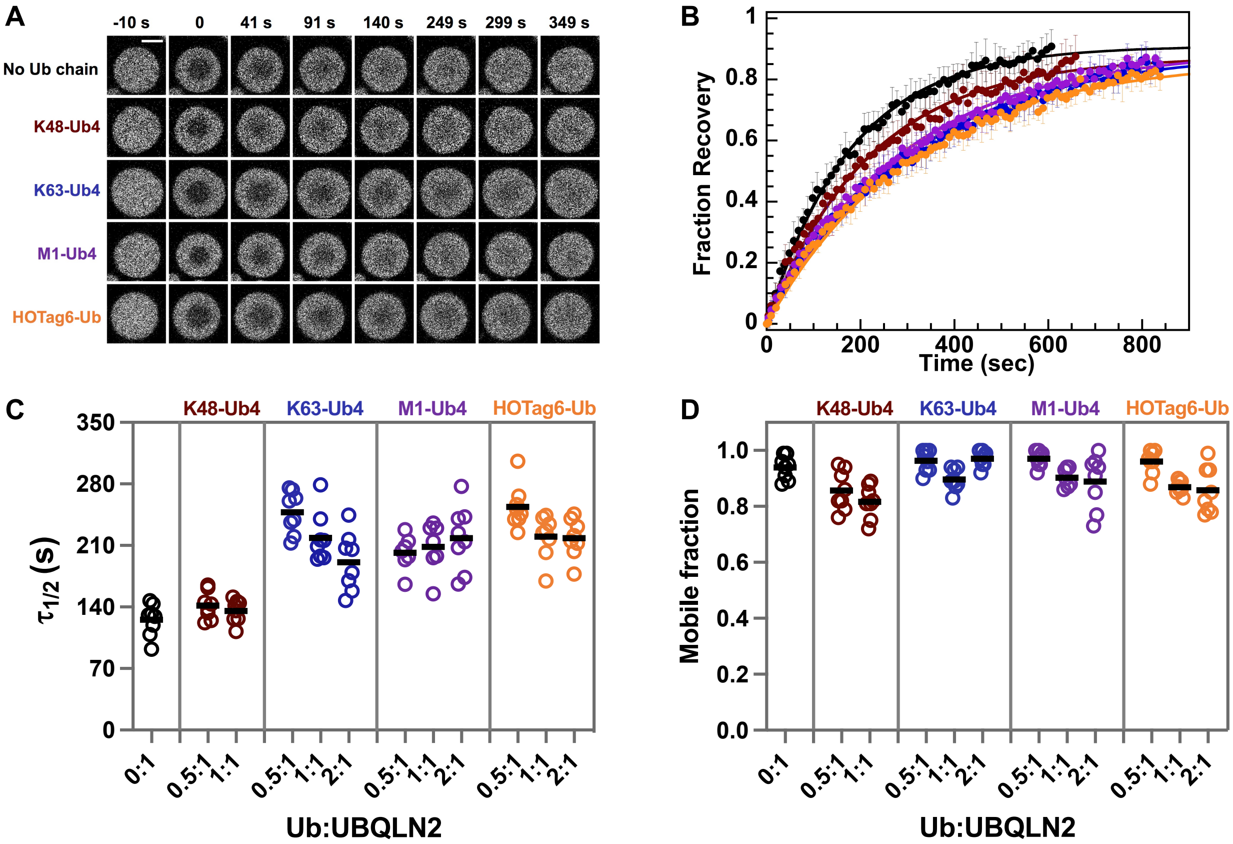


**Supplementary Figure 8.** FRAP of UBQLN2 droplets in the presence of different Ub4 chains. (A) Representative fluorescence images of partial droplet photobleaching experiments for UBQLN2 droplets in the absence or presence of 1:1 Ub:UBQLN2 molar ratio of K48-Ub4, K63-Ub4, M1-Ub4 and HOTag6-Ub to UBQLN2. Scale bar is 5 µm. (B) Normalized fluorescence intensity over time for droplets in the absence or presence of 1:1 molar ratio of UBQLN2:chain. Dots indicate the average data points from 7-8 separate droplets. Thick lines indicate a single exponential fit to the data. Error bars represent the SD. (C) Half-times of fluorescence recovery and (D) mobile fractions of UBQLN2 in the absence or presence of different molar ratios of K48-Ub4, K63-Ub4, M1-Ub4 and HOTag6-Ub to UBQLN2. Black lines represent the mean.

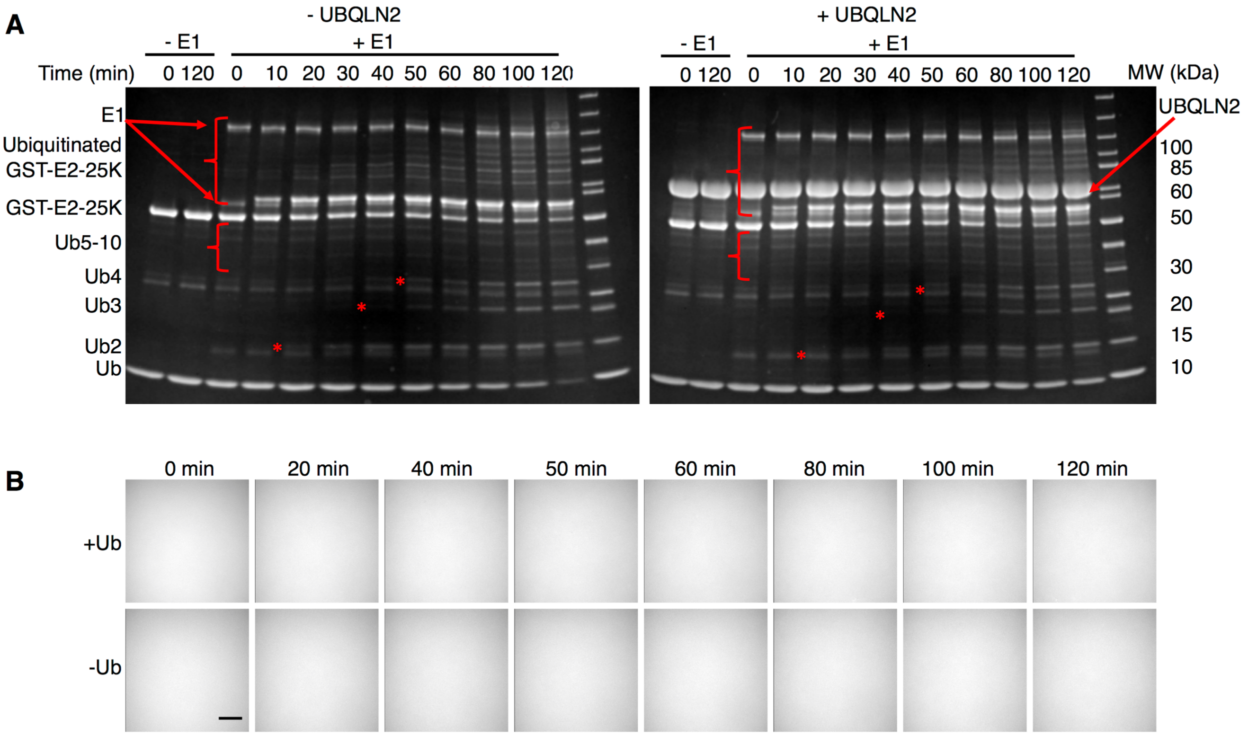


**Supplementary Figure 9.** (A) SDS-PAGE gels monitoring the formation of K48-linked polyUb chains and possibly ubiquitinated GST-E2-25K over time, as indicated by asterisks and curly brackets. Conditions: ± 50 µM Ub and UBQLN2, 30 nM Dylight 650-labeled UBQLN2, 1 µM mE1, 10 µM GST-E2-25K, in buffer containing 10 mM ATP and $MgCl_2$, 3 mM TCEP, 50 mM Tris pH 8, at 37 ºC. The K48-linked polyUb chain reactions were done in the absence (left) and presence (right) of UBQLN2. Since UBQLN2 competes with E1 and E2 to bind to Ub, the presence of UBQLN2 slowed down chain formation (indicated by comparing disappearance rates of Ub). The band intensity for GST-E2-25K decreased with time. The time-dependent appearance of species between 50-100 kDa were obstructed in the presence of UBQLN2. Since GST-E2-25K is not as efficient at making K48-linked polyUb chains as Mms2 and GST-Ubc13 are at making K63-linked chains, we carried out the experiments for 2 hours to get a comparable rate (by looking at the disappearance of Ub). (B) Time-lapse fluorescence microscopy monitoring UBQLN2 droplet formation in the presence of Ub and K48-selective ubiquitination machinery. Experimental conditions were the same as in (A). Imaging was done in solution above the coverslip. Scale bar, 5 µm.

**Table S1. $K_d$ determined from NMR and fluorescence anisotropy titration experiments.**

| | NMR-derived [a] | FA-derived (labelled 450-624) [b] | FA-derived (labelled Ub4) [b] |
|---|---|---|---|
| 450-624 with Ub | 3.38 ± 0.20 | 3.36 ± 1.17 | |
| 450-624 with K11-Ub4 | 4.63 ± 1.32 | 0.39 ± 0.08 | |
| 450-624 with K48-Ub4 | 25.34 ± 7.51 | 0.42 ± 0.25 | 2.76 ± 0.23 |
| 450-624 with K63-Ub4 | 3.29 ± 1.40 | 2.29 ± 0.55 | |
| 450-624 with M1-Ub4 | 12.83 ± 2.81 | 5.11 ± 1.30 | |
| 450-624 with HOTag6-G10-Ub | 8.83 ± 0.93 | 6.98 ± 1.72 | |

| | NMR-derived $k_{off}$ [c] |
|---|---|
| 450-624 with Ub | 861.1 ± 28.1 |
| 450-624 with K11-Ub4 | 168.1 ± 8.1 |
| 450-624 with K48-Ub4 | 232.1 ± 68.4 |
| 450-624 with K63-Ub4 | 293.3 ± 42.3 |
| 450-624 with M1-Ub4 | 495.0 ± 67.2 |
| 450-624 with HOTag6-G10-Ub | 605.2 ± 44.0 |

[a] $K_d$ was determined using TITAN analysis with a single-site binding equation using global fitting of multiple residues (see Methods and Figure S6). Errors are from jackknife analysis.

[b] $K_d$ was determined from the average $K_d$ (μM) from three separate trials. Errors are the standard deviation of the $K_d$ values.

[c] $k_{off}$ was determined using TITAN analysis with a single-site binding equation using global fitting of multiple residues (see Methods and Figure S6). Errors are from jackknife analysis.

**Table S2. Structural parameters of polyUb from SAXS data analysis.**

| Samples | $R_g$ (Å) [a] (Guinier) | $R_g$ (Å) [b] (GNOM) | Dmax (Å) (GNOM) |
|---|---|---|---|
| K48-Ub4 | 25.67 ± 0.05 | 26.19 ± 0.07 | 97 |
| K63-Ub4 | 32.40 ± 0.07 | 34.17 ± 0.14 | 133 |
| M1-Ub4 | 32.42 ± 0.03 | 34.22 ± 0.21 | 140 |
| HOTag6-G10-Ub | 37.81 ± 0.06 | 39.06 ± 0.16 | 135 |

Indicated in parentheses are methods/software used for $R_g$ analysis.

[a] The $R_g$ and errors are determined from the linear fit of ln(I(q)) vs. $q^2$.

[b] The $R_g$ and errors are determined from choosing multiple values of $D_{max}$.

**Table S3. Summary of SV-AUC experiments with UBQLN2, Ub4 chains, and UBQLN2:Ub4 complexes.**

| Samples | s*, $s_{20,w}$ [a] | $f/f_0$ [b] | Loading Concentration (μM) | MSSV [c] Calculated Concentration (μM) | MSSV Calculated Stoichiometry of Complex |
|---|---|---|---|---|---|
| UBQLN2 487-624 | 1.23, 1.25 | 1.701 | 200 | - | - |
| K48-Ub4 | 2.58, 2.64 | 1.346 | 100 | - | - |
| K63-Ub4 | 2.49, 2.55 | 1.394 | 100 | - | - |
| UBQLN2: K48-Ub4 complex [d] | 2.8, 2.84 | 1.617 | 200 : 100 UBQLN2:Ub4 | 190.0 : 98.8 UBQLN2:Ub4 | 1.9:1 |
| UBQLN2: K63-Ub4 complex [d] | 2.7, 2.7 | 1.635 | 200 : 100 UBQLN2:Ub4 | 196.7 : 95.0 UBQLN2:Ub4 | 2.1:1 |

[a] Experimental sedimentation coefficients (s*) and standard sedimentation coefficients ($s_{20,w}$) after correcting for water at 20 $^{o}$C.

[b] Frictional ratio.

[c] MSSV analysis was performed to determine protein concentrations in the complex and overall binding stoichiometry using parameters in Table S4 (see Methods).

[d] UBQLN2 here is UBQLN2 487-624, a construct previously shown to be monomeric (Dao et al., 2018).

**Table S4. Parameters for MSSV analysis.**

| Samples | $\varepsilon_k$ $\lambda$=260 ($M^{-1}$ $cm^{-1}$) | $\varepsilon_k$ $\lambda$=280 ($M^{-1}$ $cm^{-1}$) | Ratio [a] |
|---|---|---|---|
| UBQLN2 487-624 | 5555 | 5500 | 1.01 |
| K48-Ub4 | 9882 | 5960 | 1.66 |
| K63-Ub4 | 10810 | 5960 | 1.81 |

[a] $D_{norm}$ = 0.2730/0.2344 (values > 0.06 are considered good for spectral discrimination).

**Table S5. SEC-MALS-SAXS Data Collection and Analysis.**

| (*a*) Sample details | | | | |
|---|---|---|---|---|
| | 1 | 2 | 3 | 4 |
| Organism | *H. sapiens* | *H. sapiens* | *H. sapiens* | *H. sapiens* |
| Source (Catalogue No. or reference) | *Expressed in E. coli (this work)* | *Expressed in E. coli (this work)* | *Expressed in E. coli (this work)* | *Expressed in E. coli (this work)* |
| Description: sequence (including Uniprot ID + uncleaved tags), bound ligands/modifications, *etc.* | K48-Ub4 (distal Ub is K48R, all other Ub units are WT Ub) | K63-Ub4 (distal Ub is K63R, all other Ub units are WT Ub) | M1-Ub4 (linear tetraubiq.) | HOTag-$G_{10}$-Ub (see Methods) |
| Extinction coefficient ε in $M^{-1}$ $cm^{-1}$ (wavelength in nm) | 5960 (280) | 5960 (280) | 5960 (280) | 6990 (280) |
| Molecular mass *M* from chemical composition (Da) | 34,200 | 34,200 | 34,200 | 13,200 |
| For SEC-SAS, loading volume/concentration (mg $ml^{-1}$), injection volume (μl), flow rate (ml $min^{-1}$) | 7.0, 300, 0.6 | 7.0, 300, 0.6 | 7.0, 300, 0.6 | 4.5, 300, 0.6 |
| Solvent composition and source | pH 6.8 20 mM NaPhos, 0.5 mM EDTA, 0.02% $NaN_3$ | pH 6.8 20 mM NaPhos, 0.5 mM EDTA, 0.02% $NaN_3$ | pH 6.8 20 mM NaPhos, 0.5 mM EDTA, 0.02% $NaN_3$ | pH 6.8 20 mM NaPhos, 0.5 mM EDTA, 0.02% $NaN_3$ |

| (*b*) SAS data collection parameters | |
|---|---|
| Source, instrument and description or reference | BioCAT facility at the Advanced Photon Source beamline 18ID with Pilatus3 X 1M (Dectris) detector |
| Wavelength (Å) | 1.033 |
| Beam geometry (size, sample-to-detector distance) | 150 (h) x 25 (v) focused at the detector |
| Camera Length (m) | 3.658 |
| *q*-measurement range ($Å^{-1}$) | 0.003-0.35 |
| Absolute scaling method | Glassy Carbon, NIST SRM 3600 |
| Basis for normalization to constant counts | To transmitted intensity by beam-stop counter |
| Method for monitoring radiation damage, X-ray dose where relevant | Automated frame-by-frame comparison of relevant regions using CORMAP (Franke et al., 2015) implemented in BioXTAS RAW |
| Exposure time, number of exposures | 0.5 s exposure time with a 2 s total exposure period (0.5 s on, 1.5 s off) of entire SEC elution |
| Sample configuration including path length and flow rate where relevant | SEC-MALS-SAXS. Size separation used a Superdex 200 10/300 Increase column and a |

| | |
|---|---|
| | 1260 Infinity II HPLC (Agilent Technologies). UV data was measured in the Agilent, and<br>MALS-DLS-RI data by DAWN HELEOS-II (17 MALS + 1 DLS channels) and Optilab T-rEX (RI) instruments (Wyatt Technology). SAXS data was measured in a sheath-flow cell (Kirby et al., 2016), effective path length 0.542 mm. |
| Sample temperature (ºC) | 20 |

(*c*) Software employed for SAXS data reduction, analysis and interpretation

| | |
|---|---|
| SAXS data reduction | Radial averaging; frame comparison, averaging, and subtraction done using BioXTAS RAW 2.1.1 (Hopkins et al., 2017) |
| Calculation of ε from sequence | ProtParam (ExPASy) |
| Basic analyses: Guinier, M.W., *P*(*r*), scattering particle volume (*e.g.* Porod volume $V_P$ or volume of correlation $V_c$) | Guinier fit and M.W. using BioXTAS RAW, P(r) function using GNOM (Svergun, 1992). RAW uses MoW and Vc M.W. methods (Piiadov et al., 2019; Rambo and Tainer, 2013) |
| MALS-DLS-RI analysis | Astra 7 (Wyatt) |

(*d*) Structural parameters

| Guinier Analysis | K48-Ub4 | K63-Ub4 | M1-Ub4 | HOTag-Ub |
|---|---|---|---|---|
| *I*(0) ($cm^{-1}$) | 0.0306 ± 0.00003 | 0.0326 ± 0.00004 | 0.02832 ± 0.00004 | 0.02688 ± 0.00006 |
| $R_g$ (Å) | 25.67 ± 0.05 | 32.40 ± 0.07 | 32.42 ± 0.03 | 37.81 ± 0.06 |
| *q*-range (Å$^{-1}$) | 0.003 – 0.05127 | 0.003 – 0.04013 | 0.003 – 0.03984 | 0.00443 – 0.03556 |
| Quality-of-fit parameter (with definition) | 0.9943 ($r^2$) | 0.9946 ($r^2$) | 0.9920 ($r^2$) | 0.9966 ($r^2$) |
| *M* from MALS (kDa) | 33.7 | 34.5 | 33.9 | 53.4 |
| *P*(*r*) analysis | K48-Ub4 | K63-Ub4 | M1-Ub4 | HOTag-Ub |
| *I*(0) ($cm^{-1}$) | 0.0306 | 0.0352 | 0.0286 | 0.0271 |
| $R_g$ (Å) | 26.17 | 34.04 | 34.22 | 39.06 |
| $d_{max}$ (Å) | 97 | 135 | 140 | 135 |
| *q*-range (Å$^{-1}$) | 0.003 – 0.3491 | 0.003 – 0.3491 | 0.003 – 0.03491 | 0.003 – 0.3491 |
| Quality-of-fit parameter (with definition) | 1.0441 (chi2) | 1.0522 (chi2) | 1.0373 (chi2) | 1.0075 (chi2) |

### Supplementary Movie

**Movie S1. UBQLN2 undergoes phase separation in the presence of Ub and K63-selective ubiquitination machinery, related to Fig. 6.**

Time-lapse fluorescence microscopy monitoring UBQLN2 droplet formation in the presence of Ub and K63-selective ubiquitination machinery. Imaging was done in solution above the coverslip. Movie is about 31 minutes in real time. Initial experimental conditions: 50 μM Ub and UBQLN2, 30 nM Dylight 650-labeled UBQLN2, 1 μM mE1, 2 μM His-Mms2, 4 μM GST-Ubc13, 10 mM ATP and $MgCl_2$, 3 mM TCEP, 50 mM Tris pH8, 37 ºC. As time goes on, the amount of Ub in solution decreases while the amounts of different K63-linked polyUb chains increase.
